# Constructing Gene Regulatory Networks using Epigenetic Data

**DOI:** 10.1101/2020.10.19.345827

**Authors:** Abhijeet Rajendra Sonawane, Dawn L. DeMeo, John Quackenbush, Kimberly Glass

## Abstract

The biological processes that drive cellular function can be represented by a complex network of interactions between regulators (transcription factors) and their targets (genes). A cell’s epigenetic state plays an important role in mediating these interactions, primarily by influencing chromatin accessibility. However, effectively leveraging epigenetic information when constructing regulatory networks remains a challenge. We developed SPIDER, which incorporates epigenetic information (DNase-Seq) into a message passing framework in order to estimate gene regulatory networks. We validated SPIDER’s predictions using ChIP-Seq data from ENCODE and found that SPIDER networks were more accurate than other publicly available, epigenetically informed regulatory networks as well as networks based on methods that leverage epigenetic data to predict transcription factor binding sites. SPIDER was also able to improve the detection of cell line specific regulatory interactions. Notably, SPIDER can recover ChIP-seq verified transcription factor binding events in the regulatory regions of genes that do not have a corresponding sequence motif. Constructing biologically interpretable, epigenetically informed networks using SPIDER will allow us to better understand gene regulation as well as aid in the identification of cell-specific drivers and biomarkers of cellular phenotypes.

## Introduction

The regulatory program of a cell can be defined by a complex set of interactions that occur when transcription factors (TFs) bind and recruit the transcriptional machinery to the regulatory regions of their target genes^1, 2^. Modeling these interactions in gene regulatory networks allows for the identification of cell-specific regulatory processes, providing insights into how cells develop, respond to environmental perturbations, and are altered by disease. However, gene regulation is a complex process. Transcription factors form protein complexes, leading to co-regulatory events even in the absence of corresponding recognition sequences. These events are mediated through various mechanisms, including DNA looping, piggyback recruitment, assisted binding, or interaction with modified histones^2^. In particular, the epigenetic state of the cell, through chromatin architecture and nucleosome positioning, plays a critical role in transcription factor binding events and, consequently, gene regulation.

Enzymatic chromatin accessibility assays like DNase-seq^3^ and ATAC-seq^4^ can identify regions of open (accessible) chromatin that are “protected” by bound proteins such as transcription factors, but these assays do not provide the identity of the factors bound within these regions. In contrast, high-throughput DNA binding experiments, such as ChIP-seq^5^, provide the genomic locations of specific TFs in a given context. However, these assays cover only a small number of TFs due to a lack of good antibodies and the cost associated with running many sequencing assays. Consequently, TFs are generally associated with genes by identifying transcription factor binding sites (TFBS) using DNA recognition sequences, called motifs^6–8^. Combining DNA accessibility (and other omic data) with motif information is commonly done to improve TFBS prediction^9–16^ and infer the potential regulatory roles of transcription factors^17, 18^.

TFBS prediction methods generally score each TF motif instance and validate performance based on ChIP-seq TF binding data. Regulatory networks based on TFBS predictions, leverage the subset of these motif instances that map within the promoter (or other regulatory) regions of genes. However, restricting the data to only regulatory regions drastically reduces the network’s predictive performance compared to that observed in genome-wide – and thereby gene location-agnostic – assessments (see **Supplemental Figure 1** and **Supplemental Section S1**). This indicates that TFBS information alone – even that which incorporates chromatin state and other genomic data – is insufficient for reconstructing an accurate regulatory network. This may be due to the fact that TFBS predictions do not generally incorporate any of the higher-order, co-regulatory mechanisms that are mediated by epigenetic factors.

Here we propose a method to reconstruct gene regulatory networks based on information exchange between epigenetically accessible motifs. SPIDER (Seeding PANDA Interactions to Derive Epigenetic Regulation) integrates TF motif locations with epigenetic data and then applies a message passing algorithm (PANDA) to construct gene regulatory networks. We applied SPIDER to DNase-seq data for six human cell lines and evaluated the predicted networks using independently derived ChIP-seq data. We find that SPIDER significantly outperforms other methods. Importantly, we also show that SPIDER’s unique approach of melding epigenetic data with message passing allows for the detection of indirect regulatory events. An implementation of SPIDER is available at: https://github.com/kimberlyglass/spider.

## Results

### SPIDER: Seeding PANDA Interactions to Derive Epigenetic Regulation

SPIDER is based on identifying transcription factor (TF) motifs found in accessible chromatin regions, using this information to identify an initial “seed” network, and then applying message passing to harmonize connections across all the transcription factors and genes (**Figure 1a**). SPIDER repurposes the message passing approach implemented in PANDA, a multi-omic network reconstruction algorithm based on affinitypropagation.^19^ PANDA constructs regulatory networks by integrating transcription factor motif information with protein interaction and gene co-expression data. PANDA has been widely applied a wide range of biological problems, including the study of human diseases^20–22^, tissues^23, 24^, and cell lines^25–27^. While PANDA has been extremely successful, it does not explicitly incorporate epigenetic data. This means that in practice the input TF-gene seed network used by PANDA often assumes that all motif sites on the genome are equally accessible.

**Figure 1:**
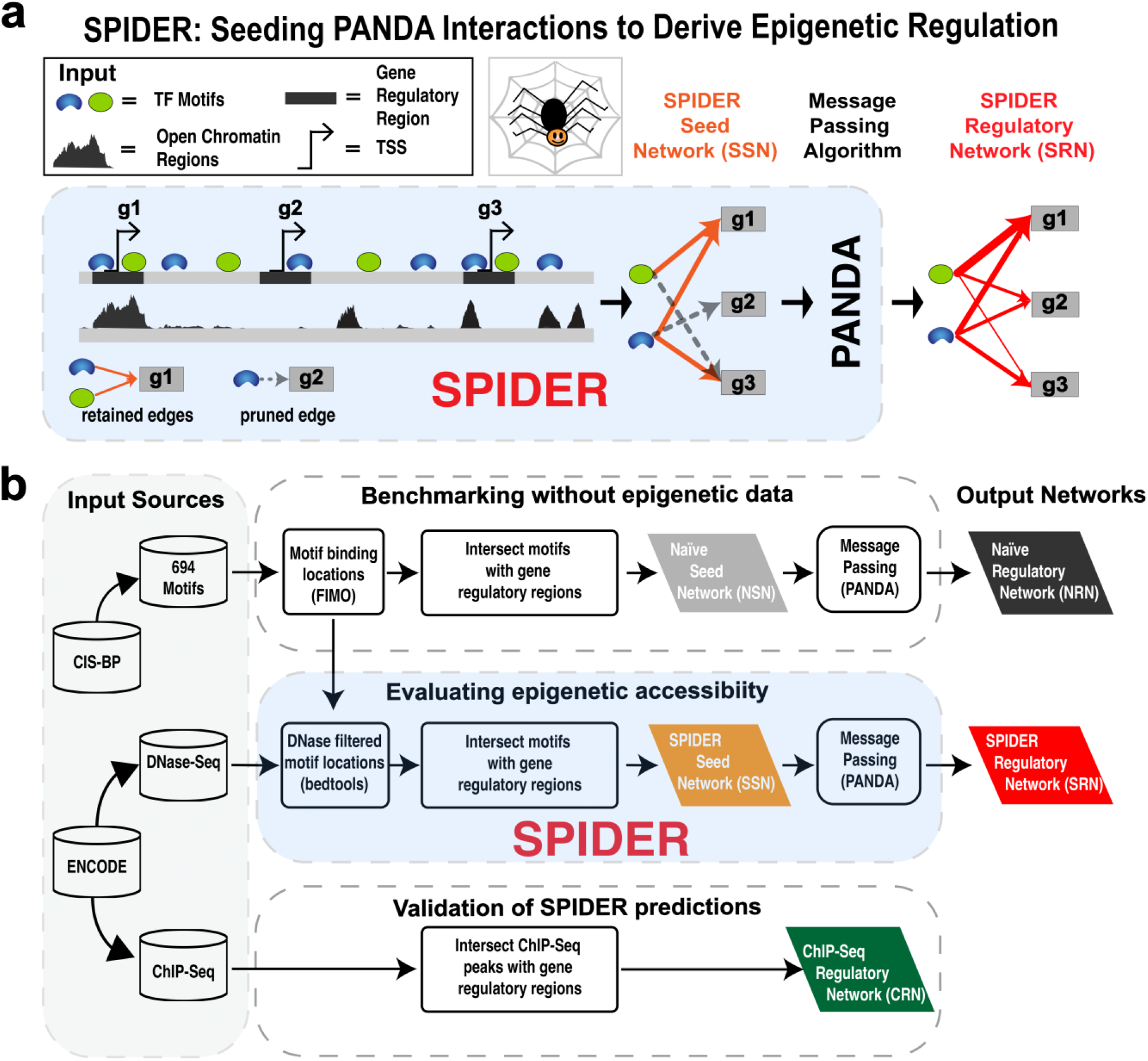
Overview of SPIDER and evaluation pipeline. **a.** Schematic of the SPIDER network reconstruction approach. **b**. Overview of the pipeline we used to evaluate SPIDER, including input data sources, key algorithmic steps, and output networks assessed.

The input to SPIDER includes the location of (1) TF motifs, defined by position weight matrices mapped onto the DNA^28^, (2) open chromatin regions, based on epigenetic data, and (3) gene regulatory regions, which can be defined based on proximity to transcriptional start sites. SPIDER first constructs a “seed” regulatory network between transcription factors and target genes by intersecting motif locations with open chromatin and gene regulatory regions. Next, the weights of edges in this initial network are degree-normalized to emphasize connections to high degree TFs and genes (see **Supplemental Section S2.1**); by definition, these TFs and genes are associated with more open chromatin regions and are therefore more likely to be active players in the regulatory process. This initial network is then run through PANDA’s message passing algorithm to optimize the network structure given the input data. It should be noted that, although PANDA generally incorporates proteinprotein interaction (PPI) and gene co-expression data, these data are not required for the message passing procedure. Because we wanted to understand how chromatin structure information influences regulatory network reconstruction, we chose to exclude PPI and expression data when testing SPIDER. The output of SPIDER is a bipartite, complete network with weighted edges that represent the likelihood of a regulatory relationship between a TF and its target genes. For a more detailed description of the SPIDER method, see **Supplemental Section S2.1**.

We tested SPIDER using data for six human cell lines (**Table 1**). In particular, for the input to SPIDER we used (1) TF motif data derived from mapping transcription factors motifs from Cis-BP^29^ to the hg19 genome using FIMO^28^, (2) open chromatin regions defined in narrow-Peak DNase-seq data files from ENCODE, and (3) regulatory regions, defined as 2kb windows centered around the transcriptional start sites of genes based on RefSeq annotations^30^ (**Figure 1b**). For each cell line we estimated two epigenetically-informed networks: the initial “seed” network constructed from integrating TF binding sites with open chromatin and gene regulatory regions (orange parallelogram in **Figure 1b**) and a final SPIDER regulatory network estimated from performing message passing on the seed network (red parallelogram). We also estimated two reference networks common to all six cell lines: an initial “naïve” network consisting of TF-gene associations based only on the intersection of motif locations with regulatory regions promoters (gray parallelograms), and a final naïve network derived by applying message passing to refine the initial naïve network (black parallelograms). These networks included regulatory associations for 687 regulating TFs and 27090 target genes. Finally, we created six “gold standard” validation networks, one for each of the six cell lines, by taking the intersection of experimental cell-specific ChIP-seq peaks with regulatory regions. It should be noted that the dimensions of these ChIP-seq networks vary based on the number of TFs assayed in each cell line (**Table 1**; **Supplemental File 1**). In total we have twenty reconstructed networks – three for each cell line (SPIDER seed network, SPIDER regulatory network, ChIP-seq regulatory network) plus the “naïve” seed and regulatory networks – as well as six “gold standard” networks based on ChIP-seq data. This set of networks allowed us to explore both the impact of including epigenetic information as well as the message passing optimization in SPIDER. For more information on our data processing and network construction pipeline, see **Supplemental Sections S2.2-S2.3**.

**Table 1:** Overview of human cell lines used in this paper together with information on each cell line, the number of unique TF motifs with corresponding ChIP-seq data that we evaluated in that cell line, and the number of DNase-seq peaks.

| Cell line | Tissue | Description | # ChIP-seq TFs | # DNase peaks |
| --- | --- | --- | --- | --- |
| A549 | Lung | Alveolar Carcinoma | 19 | 176870 |
| H1HESC | Stem cell | Male embryo stem cell | 35 | 258188 |
| HELAS3 | Cervix | Cervical adenocarcinoma | 38 | 199188 |
| HEPG2 | Liver | Hepatocellular carcinoma | 45 | 192959 |
| GM12878 | Blood | B-lymphoblastoid cells | 58 | 183953 |
| K562 | Blood | Erythrocytic leukemia | 59 | 202266 |

### SPIDER predicts accurate gene regulatory networks

To begin, we benchmarked the two naïve networks using the six ChIP-seq “gold standard” networks and evaluated their accuracy based on the Area Under the Receiver-Operator Characteristic Curve (AUC-ROC, or more simply, AUC). This provided a baseline assessment of network accuracy in the absence of epigenetic information (**Figure 2a**). Unsurprisingly, we observed very low AUC values – from ~0.57 to ~0.60 (**Figure 2b**, gray and black bars) – indicating that message passing in the absence of cell line specific epigenetic data does not improve the network accuracy. We also evaluated the SPIDER seed networks, which represent the contribution of chromatin accessibility data to network accuracy in the absence of applying message passing (**Figure 2b**, pale colored bars). Interestingly, we found only a marginal increase in AUC compared to the naïve networks. For example, the SPIDER seed networks for A549 had an AUC of 0.594, only a 3.8% increase over the naïve seed network (AUC=0.572). This indicates that the addition of epigenetic data alone does not result in accurate gene regulatory networks. For more explanation regarding these results, see **Supplemental Section S1**.

**Figure 2:**
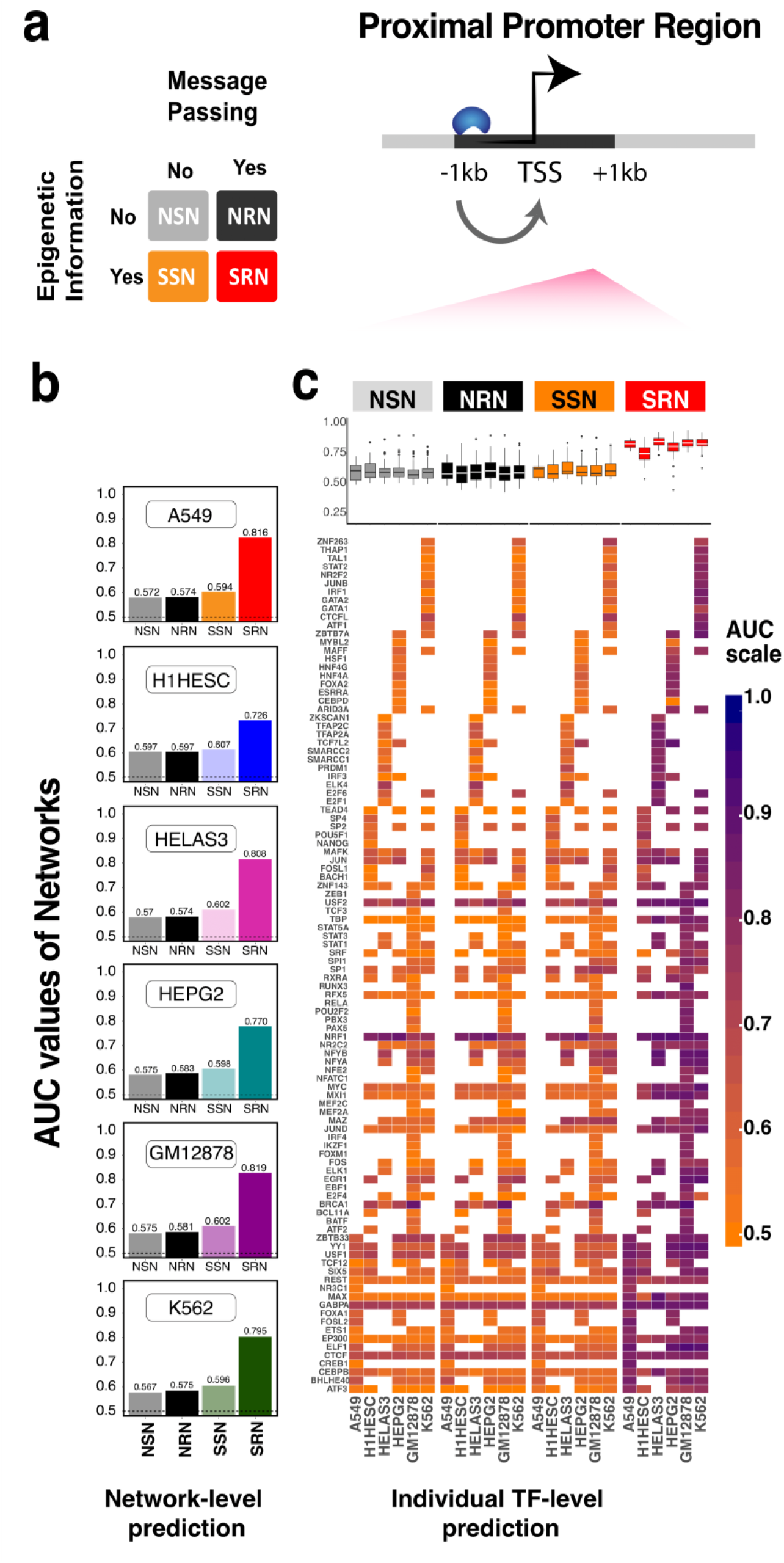
Evaluation of SPIDER predictions. **a.** For each cell line we evaluated four networks, modeled either with or without using epigenetic data, as well as with or without applying message passing. SPIDER-predicted regulatory networks represent a combination of epigenetic information and message passing. **b**. The AUC values of four types of networks evaluated in six different cell lines (based on ChIP-seq gold standards). A baseline AUC value of 0.5 is shown as a horizontal dotted line. **c**. The AUC values for individual TFs within each network. The distribution of values is shown in the top panel. Individual values are visualized in the bottom panel. See also **Supplemental File 1**, **Supplemental Table 1**, and **Supplemental Figure 2**.

We next evaluated the SPIDER predicted gene regulatory networks, which are the result of applying message passing to the epigenetically informed seed networks (**Figure 2b**, dark colored bars). We found that SPIDER regulatory networks are highly accurate, with AUC scores dramatically increased compared to both the naïve and epigenetically informed SPIDER seed networks. For example, the accuracy of the A549 network was improved by over 37% (AUC=0.816) compared the seed network. The SPIDER predicted network for GM12878 was the most accurate, with an AUC value of 0.819. This level of accuracy and overall improvement was consistent across all six cell lines (**Supplemental Table 1**). This is remarkable, especially given that adding epigenetic data to the naïve networks did little to improve accuracy. It also demonstrates the robustness of SPIDER in predicting accurate regulatory networks across a range of cell types. Importantly, these results indicate that the combination of epigenetic information and message passing, rather than either in isolation, is critical for uncovering the cellular regulatory architecture.

To ensure that these results were not driven by a handful of transcription factors, such as those with a high number of motif locations or abundant ChIP-seq peaks, we separately evaluated the accuracy of the edges emanating from each individual transcription factor (**Figure 2c**). Just as with the overall networks, we found that the AUC values for TFs were nearly always significantly higher in SPIDER networks than in the corresponding naïve and seed networks, and that this was true across all the cell lines. Of note, transcription factors with ChIP-seq data in multiple cell lines (CEBPB, CTCF, EP300, GABPA, MAX, REST in all six cell lines; ATF3, USF1, YY1, JUND, MX11, MYC, NRF1, RFX5, TBP, USF2, in five cell lines) had consistent AUC values across the cell lines. Interestingly, some of these transcription factors also have higher AUC values across all the different network types. This may be due to the fact that their corresponding motif is found in the promoter region of more genes. For example, GABPA targets 18-19% of all genes in the seed networks, whereas the average TF only targets about 6% of genes in the seed networks. Other examples include CTCF (targets ~10% of genes), NRF1 (~10%), and USF2 (~7%).

Finally, we verified that these results were not the result of class-imbalance by repeating these analyses using the Area Under the Precision-Recall Curve (AUPR) instead of AUC to evaluate network accuracy. We observed almost identical results using AUPR as we did with AUC (**Supplemental Figure 2**). Overall, these analyses demonstrate that SPIDER effectively applies message passing to an epigenetically informed seed network in order to infer accurate gene regulatory networks. Importantly, these networks represent the *in vivo* regulatory architecture observed in ChIP-seq data, both at the overall gene regulatory network level and at the individual transcription factor level.

### SPIDER predicts cell specific regulatory relationships

A cell’s regulatory network includes interactions that are specific to a given cell type or biological context as well as interactions that are shared across many cell types and support common regulatory processes^23^,^31^. Therefore, we next evaluated SPIDER’s ability to predict network edges that are cell line specific, i.e. instances wherein ChIP-seq data indicates that a TF is bound to the regulatory region of a gene in one cell line but not another. Such interactions are important in determining cellular function and may play a role in a wide range of cell-specific characteristics, including disease risk^32^.

To evaluate SPIDER’s ability to predict of cell line specific edges, we first constructed “differential networks”. Specifically, for each pair of cell lines (“A” and “B”), we subtracted (1) the input SPIDER seed networks, (2) the regulatory networks predicted by SPIDER, and (3) the ChIP-seq derived gold standard networks. It’s important to note that, since the seed networks and gold standard networks are binary, taking the difference between these networks results in three classes of edges: those specific to cell line A (difference = +1), those specific to cell line B (difference = −1), and those that are the same in both cell lines (either existing in both, or not existing in both; difference = 0) (**Figure 3a**).

**Figure 3:**
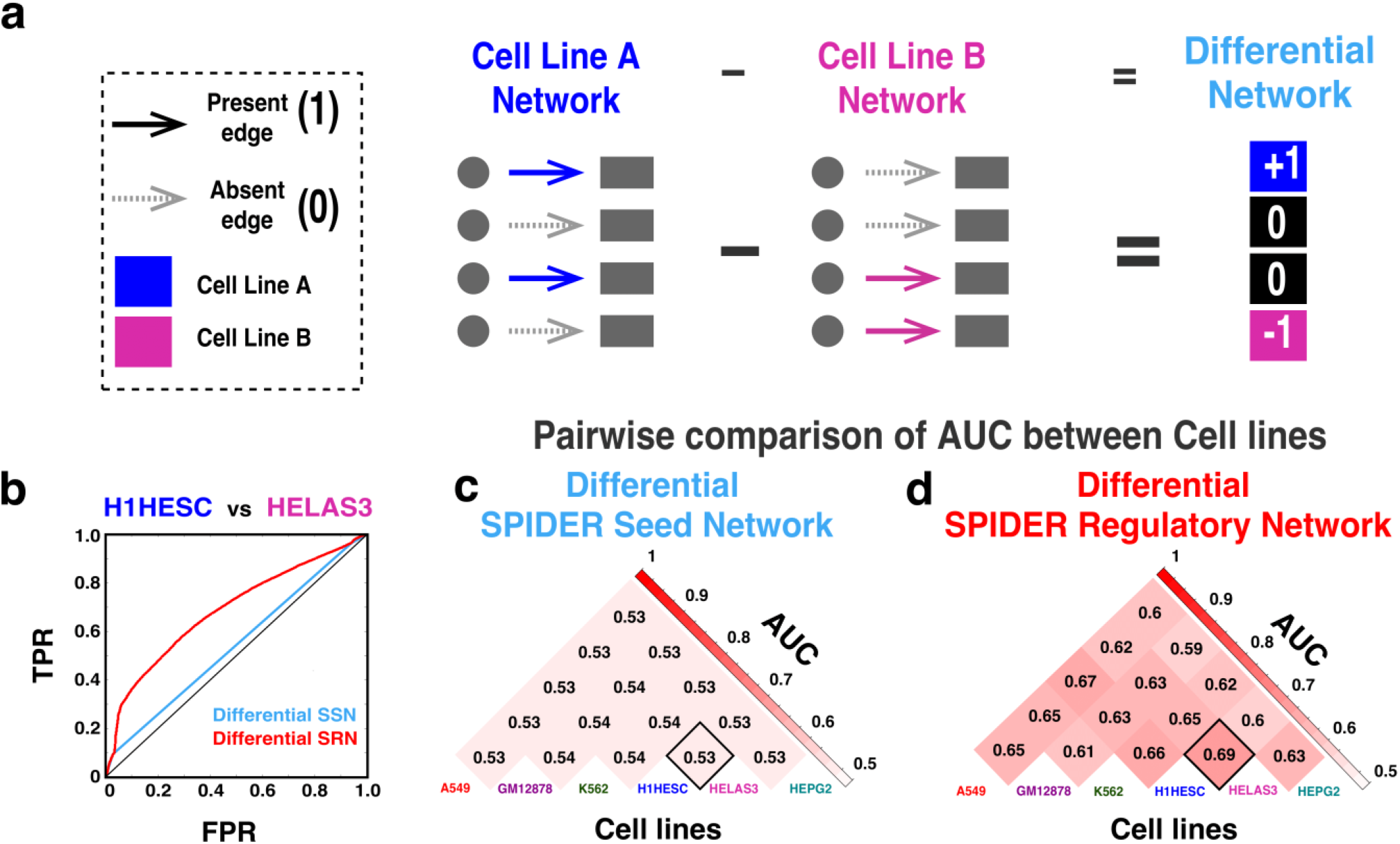
Cell-specific interactions predicted by SPIDER. **a.** Illustration of our approach to assessing cell line specific network relationships through differential network analysis. **b.** ROC curve showing the evaluation of the differential networks that were calculated by comparing the H1HESC and HELAS3 SPIDER seed networks (blue) and regulatory networks (red). **c.** AUC values of each of the pairwise comparisons of the SPIDER seed networks. **d.** AUC values of each of the pairwise comparisons of the SPIDER regulatory networks.

Next, we computed the AUC for the differential-seed and differential-regulatory networks using three-level predictors, with the differential ChIP-seq networks as our benchmark. To help illustrate and interpret this analysis, **Figure 3b** shows the ROC curve comparing one pair of cell lines: H1HESC versus HELAS3. For this pair of cell lines, the differential SPIDER regulatory network had a much higher predictive power (AUC = 0.69) compared to the differential seed network (AUC=0.53). Importantly, this curve also shows that all the three classes of edges (−1, +1, and 0) contribute to overall AUC. This can be seen by noting that the ROC curve for the differential seed network is composed of three straight line segments, one for each of the three classes of edges. The ROC for the differential SPIDER network rises above these segments and is especially pronounced for the middle segment. This segment represents edges that either existed or didn’t exist in both cell lines’ seed networks. Thus, the dramatic shift in the ROC curve indicates that the message passing portion of SPIDER is effectively removing false positives and false negatives from the epigenetically informed seed networks.

**Figures 3c-d** illustrate the AUC values for the differential networks across all pairs of cell lines. We observe that the SPIDER regulatory networks consistently predict cell line specific interactions much more accurately than the seed networks. Overall, these results indicate that message passing is enhancing the detection of cell line specific edges.

### SPIDER can be used to model distal regulatory elements

Transcription factor binding in the proximal promoter region regulates gene expression through the formation of the pre-initiation complex. Similarly, distal regulatory elements can influence the rate of gene transcription by acting as either activators or repressors^33^. Incorporating these distal regulatory factors into network models is an important step in developing a more holistic perspective on gene regulation.

One important advantage of chromatin accessibility data such as DNase-seq is the identification of enhancer regions. Although the local chromatin environment around enhancers is well studied^34–38^, less is known about which genes are targeted by these distal elements through mechanisms such as DNA looping^39–41^. However, proximity can be used to map distal regulatory elements to genes, providing a first order approximation of distal regulation. Along these lines, we modulated the definition of the regulatory region used by SPIDER to assess transcription factor binding sites located outside the proximal promoter. In particular, we defined the regulatory region of each gene as composed of two windows of 5kb each (total 10kb) located at increasing distances upstream and downstream of the TSS, up to ±100 kb. For example, **Figure 4a** shows the regulatory region of a gene as located −20 kb to −25 kb and +20 kb to +25 kb away from the TSS. For two example cell lines, GM12878 and A549, we ran SPIDER using five different definitions of potential regulatory regions: ±5-10kb, ±20-25kb, ±45-50kb, ±70-75kb, ±95-100kb. This allowed us to examine the potential impact of distal epigenetic variability on gene regulation at multiple distances; the width of these distal regions was selected such that the density of the SPIDER seed information was similar to our proximal promoter analysis. We benchmarked the results from SPIDER to their corresponding ChIP-seq derived gold-standards, also constructed based on these same regulatory regions (**Figure 4b**; **Supplemental Table 2**). The AUC values for the regulatory predictions made using these alternate windows showed little variation, indicating that SPIDER can be used to predict transcription factor binding sites outside of the proximal promoter. Interestingly, the prediction accuracy for these distal regulatory elements was even slightly higher than those obtained using proximal promoter regions.

**Figure 4:**
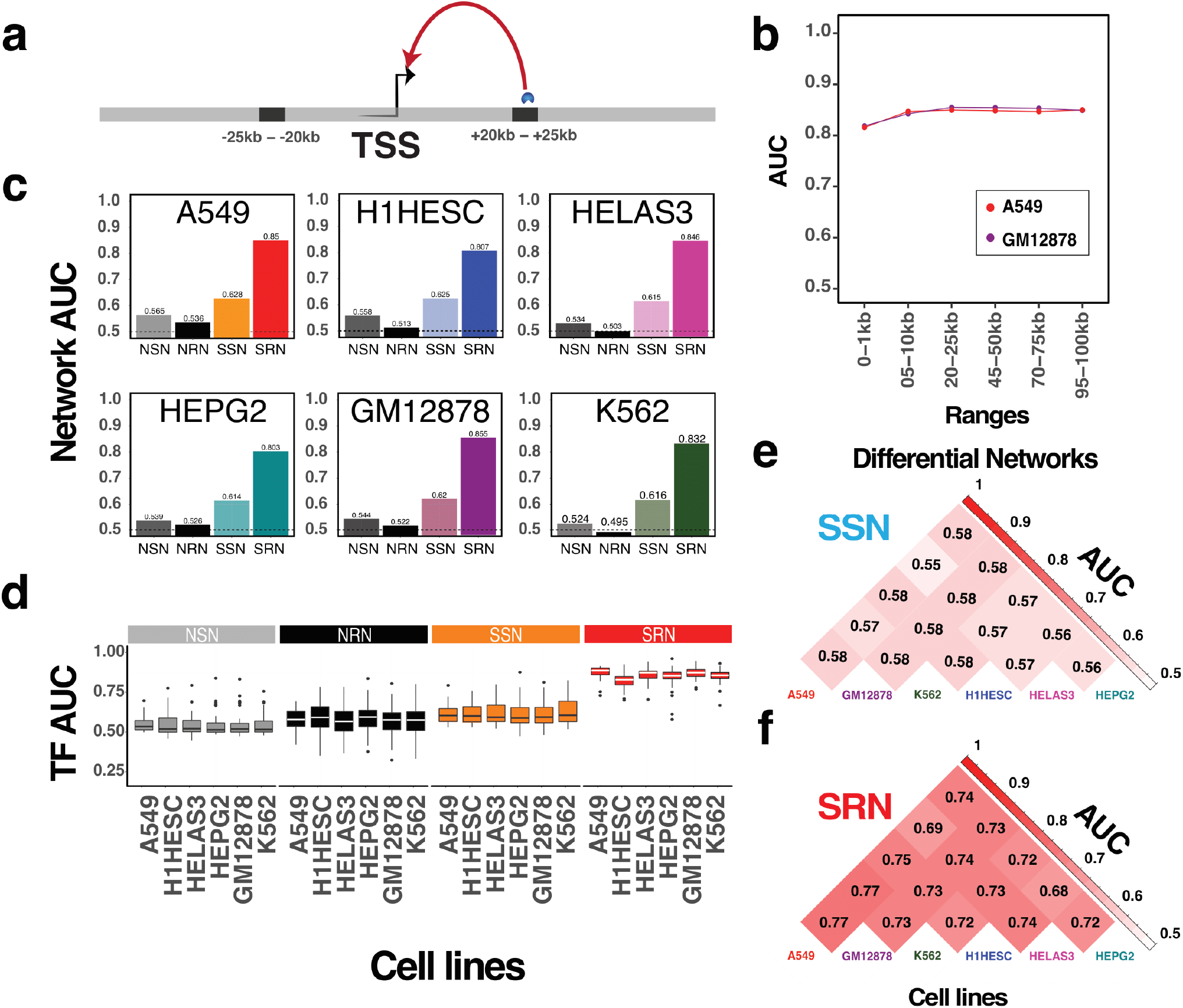
Distal regulatory events predicted by SPIDER. **a.** An example set of regions (±20-25 kb from the TSS) used to define distal regulatory windows around a gene. **b.** The AUC values for SPIDER predictions across sets of regulatory windows at increasing distance from the transcriptional start site. Two example cell lines (GM12878 and A549) are shown. **c.** The AUC values for SPIDER predictions, as well as predictions made without epigenetic information or message passing, for six cell lines using the ±20-25 kb regulatory window. **d.** Distribution of AUC values for individual TFs**. e-f.** Differential analysis demonstrates that SPIDER detects cell line specific interactions in distal regulatory windows. See also **Supplemental Table 2**, **Supplemental Table 3**, and **Supplemental Figure 3**.

Given these results, we selected a single window, ±20-25kb, to investigate in more detail. We systematically evaluated all six cell lines and found that SPIDER accurately predicts distal ChIP-seq binding events. Importantly, the accuracy of SPIDER predictions was significantly higher than either those made based on the epigenetically informed “seed” data or those based on a “naïve” mapping. This is true both overall (**Figure 4c**; **Supplemental Table 3**) as well as for individual TFs (**Figure 4d**). For example, AUC for distal regulatory interactions in A549 from SPIDER (AUC = 0.850) was over 35% higher than the seed interactions (AUC = 0.628). As in our previous analysis, we obtained very similar results when computing the AUPR instead of the AUC (**Supplemental Figure 3**). We also evaluated the cell line specificity of these distal regulatory interactions and found that the predictions made by SPIDER were highly cell-line specific and more specific than the information used to seed the algorithm (**Figure 4e-f**). These findings are consistent with the notion that biological processes specific to individual cell types or tissues are more likely to be driven by distal regulatory elements, such as enhancers, while common “housekeeping” processes tend to be regulated by promoters^42,43^.

### SPIDER outperforms other prediction algorithms

Our results demonstrate that SPIDER can effectively leverage epigenetic information to estimate highly accurate and cell line specific regulatory networks. However, there are a number of other computational tools and resources in the literature that provide epigenetically informed networks or genome-wide transcription factor binding site predictions. We benchmarked the performance of SPIDER with several of these predictive frameworks. In particular, we downloaded publicly available genome-wide transcription factor binding locations predicted by CENTIPEDE^12^ and PIQ (protein interaction quantitation)^14^, and created a set of associated gene regulatory networks by intersecting these locations with gene regulatory regions using the same pipeline we applied to build the SPIDER seed networks. We also downloaded the gene regulatory networks associated with two publications: Neph et. al.^44^ and Marbach et. al.^45^. Finally, we applied TEPIC^13^ to the same DNase data we used to test SPIDER. Additional details regarding how we processed these data is included in **Supplemental Section S3**.

It is important to note that the data reported by each of these sources varies. For example, it is possible to obtain continuous scores for the predictions made by CENTIPEDE, PIQ, and TEPIC, but the networks provided with the Neph et. al. and Marbach et. al. publications are unweighted. Therefore, to ensure a fair comparison across the sources, we converted all networks (including those estimated by SPIDER) into unweighted graphs using thresholding (see **Supplemental Section S3**). The cell lines and transcription factors included in each source also differ. Therefore, to gauge the accuracy of network predictions across these sources, we calculated a series of AUC scores by comparing the targeting profile of each transcription factor in a given cell line network with its targeting profile based on cell line specific ChIP-seq data. This resulted in a series of AUC scores associated with each method. The number of cell lines, transcription factors, and tests performed for each source is reported in **Supplemental Table 4**. The distribution of the calculated AUC values across all tests is shown in **Figure 5a**.

The networks derived from the genome-wide transcription-factor binding prediction algorithms were only marginally better than random chance; the mean AUC across all tests was only 0.516 for CENTIPEDE and 0.556 for PIQ. This is despite the fact that both of these algorithms do an outstanding job of predicting transcription factor binding at the genome-wide level (**Supplemental Figure 4a**; see **also Supplemental Section S1** and **Supplemental Figure 1**). The networks reported in Neph et. al. and Marbach et. al. were also not very accurate, with mean AUCs (based on comparison to ChIP-seq data) across all tests of 0.518 and 0.559, respectively. This is likely due to the fact that these networks were derived using similar techniques as the ones we modeled based on CENTIPEDE and PIQ. The networks predicted by TEPIC were overall more accurate than the other sources (mean AUC = 0.582) but were still less accurate than those predicted by SPIDER (mean AUC = 0.695). We note that these results were not greatly impacted by considering the scores made by the algorithm in lieu of thresholding (**Supplemental Figure 4b**) and were similar for distal regulatory interactions (**Supplemental Figure 4d-e**). As in the case of our other analyses, these results were also very similar when using the AUPR instead of the AUC to measure accuracy (**Supplemental Figure 4c&f**).

To better understand why SPIDER outperformed other methods/sources, we analyzed the components that lead to the calculation of the AUC (**Figure 5b**), namely: (1) the True Positive Rate (TPR), or instances where a TF is predicted to be regulating a gene and that gene has a ChIP-seq peak for the TF in its regulatory region; (2) the True Negative Rate (TNR), or instances where a TF is predicted to be absent and there is no ChIP-seq peak; (3) the False Positive Rates (FPR), or instances where a TF is predicted to be regulating a gene, but that prediction is not supported by ChIP-seq; and (4) the False Negative Rate (FNR), or instances where TF is predicted to be absent, but a ChIP-seq peak exists in the regulatory region of the gene (for more details, see **Supplemental Section S3**). Visualizing these rates (**Figure 5b**) revealed that although other methods generally excel at detecting true negatives, this is at the cost of greatly reducing the number of true positives, ultimately leaving to a very high false negative rate and poor overall accuracy. On the other hand, although the networks predicted by SPIDER had a slightly lower true negative rate compared to networks associated with the other sources, they also included many true positive events, which were largely missed by the other methods. This is due to the fact that, unlike previous approaches, SPIDER does not require that a TF motif is present in the promoter region of a gene in order to predict a regulatory interaction between that transcription factor and gene. Rather, an interaction between a transcription factor and a gene can be learned through SPIDER’s message passing process, which assesses the likelihood of each edge based on the overall structure of the network. Biologically, these learned relationships may represent transcription factor regulatory mechanisms that are not captured by DNA sequence^46^, such as the recruitment of cofactors^47, 48^.

### SPIDER predicts biologically relevant hidden interactions

SPIDER estimates accurate networks by simultaneously predicting two classes of regulatory relationships: those that have initial evidence based on the presence of a TF motif in the regulatory region of a gene, as well as those without evidence from TF motif data but which are instead only supported by the local structure of the regulatory network (and potentially modulated by regulatory mechanisms not encoded in the DNA sequence). We evaluated the biological significance of this second class of edges, i.e. edges which were not present in SPIDER’s seed network but had a high edge weight in SPIDER’s predicted regulatory network. For demonstration, we focused on the SPIDER seed and regulatory networks for the A549 (lung cancer) cell line. Key results for all six cell lines are included in **Supplemental Tables 5-6**.

To begin, we selected TF-gene relationships that were absent in the SPIDER seed network (i.e. true and false negative edges, or those with no evidence of TF-gene regulation based on intersecting motif data with open chromatin and gene regulatory regions; **Figure 6a**) and plotted the distribution of their weight in the predicted SPIDER regulatory network (**Figure 6b**). We then selected the subset of these relationships with the highest weights for further analysis, using FDR < 0.05 as our cutoff (see **Supplemental Section S4**); these edges are those that were absent in the epigenetically informed seed network but which were subsequently predicted after running SPIDER.

**Figure 5:**
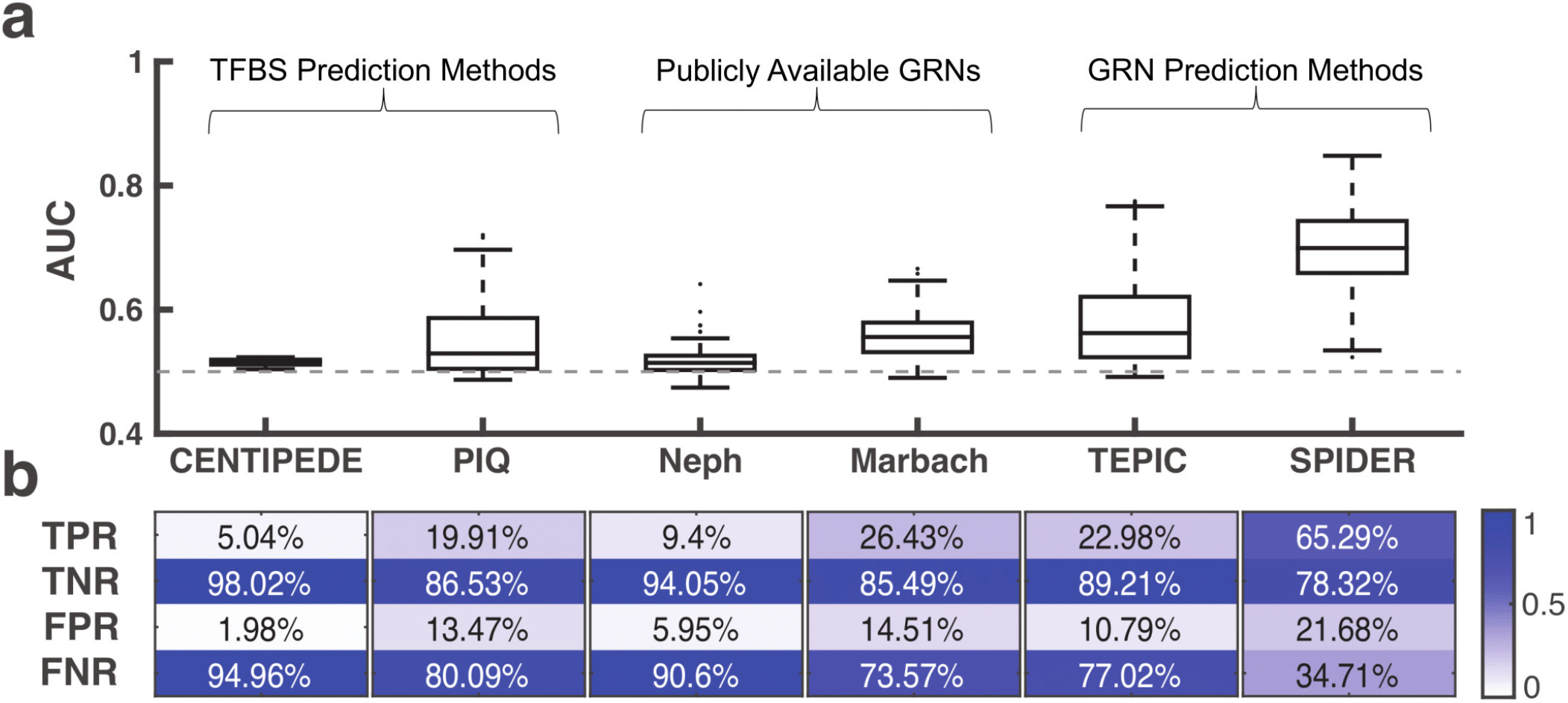
Comparison of SPIDER with other sources. **a.** An assessment of the regulatory predictions made by the networks associated with various data sources. The distribution of AUC values across all the tests performed (TF / cell line pairs) is shown. **b.** Heatmaps illustrating the median true positive rate (TPR), true negative rate (TNR), false positive rate (FPR), and false negative rate (FNR) across the tests. See also **Supplemental Table 4** and **Supplemental Figure 4**.

Next, we determined the genes targeted by these edges (**Figure 6c**). Among the genes associated with the most edges are several that are important for lung cancer, including *PDE4D* (Phosphodiesterase-4), *ZBTB20*, and *TGIF1. PDE4D* is known to promote proliferation and angiogenesis in lung cancer under hypoxia and is a potential therapeutic target for lung cancer therapy^49^. *PDE4D* is also involved in apoptosis, growth, and proliferation in lung cancer cells^50,51^, and promotes Epithelial-Mesenchymal Transition (EMT) in A549 cell lines^52^. Similarly, *ZBTB20,* a member of the POK family of transcriptional repressors, is up-regulated in lung cancer compared to adjacent normal tissue through transcriptional repression of *FOXO1*^53^. Finally, *TGIF1* knockdown inhibits the growth and the migration of non-small cell lung cancer cells^54^ and is dysregulated in several types of cancer. Assessment of *TGIF* expression has shown that silencing *TGIF* attenuates the tumorigenicity of A549 cells^55^. These results suggest that SPIDER networks can identify biologically relevant genes that may be regulated in a context-specific manner despite a lack TF motif evidence.

**Figure 6:**
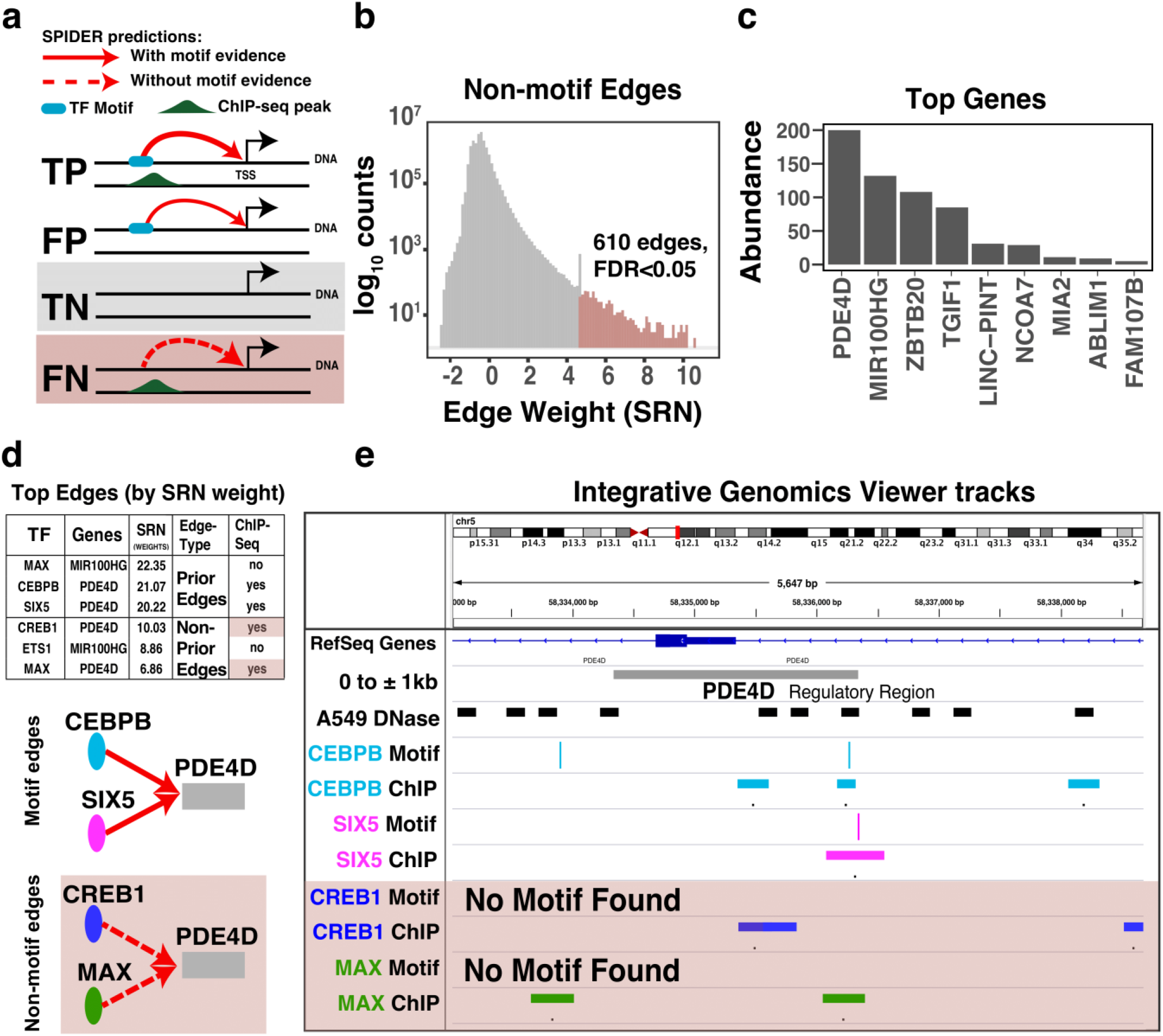
SPIDER Identification of hidden regulatory relationships. **a.** Schematic showing the relationship between true positives (TP), false positives (FP), true negatives (TN), and false negatives (FN) in the SPIDER seed network, as well as potential SPIDER predictions (red lines). **b**. Distribution of the SPIDER-predicted edge weights in the A549 network for the subset of edges that were absent in the A549 SPIDER seed network (TN or FN edges; see panel a). Of these, top-weight edges that have a significant (FDR < 0.05) weight in the SPIDER-predicted networks are shown in light red. **c.** The number of times a gene is targeted by one of the top-weight edges shown in panel b. **d.** A table showing the three top-weight edges predicted by SPIDER that originate from TFs with ChIP-seq data. Edges validated by ChIP-seq are illustrated below the table. **e.** Integrative Genomics Viewer (IGV) tracks showing DNase hypersensitivity regions, motif predictions, and ChIP-seq data in the PDE4D promoter region. Motif, DNase, and ChIP-seq peaks exist for CEBPB and SIX5. However, although only DNase and ChIP-seq peaks can be seen for CREB1 and MAX (but no corresponding motif), SPIDER recovered these regulatory relationships. See also **Supplemental Table 5** and **Supplemental Table 6**.

Finally, we investigated SPIDER-predicted transcriptional regulation, focusing on transcription factors with ChIP-seq data. In particular, we selected the three top-weight edges with motif evidence (i.e. true or false positives in the SPIDER seed network) as well as the three top-weight edges that do not have any corresponding motif evidence (i.e. true or false negatives in the SPIDER seed network). Four of these top-weight edges, including two with motif evidence and two without, targeted *PDE4D* (**Figure 6d**). To better understand these SPIDER predictions, we visualized DNase-seq, ChIP-seq, and motif locations in the regulatory region of *PDE4D* (**Figure 6e**). For CEBPB and SIX5 we find ChIP-seq peaks aligned with their corresponding motif and a DNase-seq peak. CREB1 and MAX also have ChIP-seq peaks aligned with DNase-seq peaks but no corresponding predicted motif, meaning that their potential role in regulating *PDE4D* would have been missed using other common approaches. Interestingly, CEBP proteins can mediate the binding of cAMP proteins, such as CREB1, to gene promoters^56^, suggesting that recruitment of CREB1 to *PDE4D* may have been facilitated by CEBPB – a cofactor mechanism which was likely captured by SPIDER’s message passing procedure.

## Discussion

In this manuscript we present SPIDER, a framework to predict robust, accurate, and epigenetically informed gene regulatory networks. SPIDER works by applying a message passing approach that emphasizes similarities in transcription factor targeting patterns across an initial (or seed) set of epigenetic-informed regulatory relationships. SPIDER not only predicts accurate overall networks, it also reliably estimates cell-line specific and individual transcription factor level regulatory information. Importantly, SPIDER-predicted networks are significantly more accurate than the regulatory information used to seed the algorithm as well as networks constructed without epigenetic data. This indicates that both message passing and epigenetic information are critical to SPIDER’s success.

We compared the performance of SPIDER-predicted networks to that of published gene regulatory networks and networks constructed using the results of transcription factor binding site prediction algorithms, all of which leverage similar input data. SPIDER predictions were significantly more accurate than those made by these other approaches. Other methods often require that a transcription factor’s motif is present in the regulatory region of a gene in order to assign an edge between that transcription factor and gene. Our analysis demonstrates that this leads to a high number of false negatives (missing edges). In contrast, the message passing procedure employed by SPIDER allows new TF-gene relationships to be inferred, even when a transcription factor’s motif is not present in the regulatory region of a gene.

SPIDER’s strength lies in its ability to reduce false negatives while retaining a high true positive rate, i.e. recovering missing edges without introducing a large number of false edges. SPIDER’s ability to detect and effectively enhance “hidden” interactions not only increases the accuracy of the predicted networks, it also allows for the identification of biologically important regulatory information. For example, when we investigated edges that had no supporting evidence from our motif scan, but were predicted by SPIDER, we identified PDE4D as a key target in the A549 network. PDE4D plays an important role in apoptosis, growth, and proliferation in lung cancer.

SPIDER is a highly versatile algorithm. In this manuscript, we primarily focused on modeling gene regulatory networks based on promoter regions. However, when we applied the approach to predict transcription factor binding in regions that are distally located from the transcriptional start site, we observed a very high level of accuracy. Like our promoter-based analyses, SPIDER predictions in distal regions were cell-type specific and highly accurate across transcription factors. While we recognize that the ChIP-seq data we used for validation only provides information on TF binding, and not on gene regulation, our ability to accurately predict cell-specific TF binding outside of promoter regions suggests that SPIDER can be used in modeling distal regulatory mechanisms mediated by enhancers or three-dimensional chromatin structure.

It should also be noted that we used only DNase hypersensitivity as a marker of open chromatin. This was done to facilitate the comparison of SPIDER with the extensive literature that leverages DNase data to predict transcription factor binding. However, the algorithm could easily be used with data from other epigenetic marks of open chromatin, such as ATAC-sequencing data or ChIP-sequencing of histones – this is a key future direction of our work. Finally, since SPIDER builds on the message passing framework used in the PANDA reconstruction algorithm, it has the potential to be extended to incorporate other sources of regulatory information, including protein-protein interaction and gene expression data.

SPIDER provides a principled way to use open chromatin data to gain a comprehensive understanding of the cellular transcriptional regulatory architecture. The algorithm’s unique application of message passing to highlight structures in a seed network gives it a distinct advantage compared to other methods and illustrates the importance of considering the overall regulatory context when predicting transcription factor targeting. SPIDER predicts biologically interpretable, context-specific, and epigenetically informed gene regulatory networks. Ultimately, we believe SPIDER networks will facilitate a more comprehensive understanding of regulatory processes that define health and disease.

## Data availability

An implementation of SPIDER is available at: https://github.com/kimberlyglass/spider. The input data and output networks analyzed in this manuscript can also be directly downloaded from: https://sites.google.com/a/channing.harvard.edu/kimberlyglass/tools/spider.

## Acknowledgements

We wish to thank Marieke Kuijjer, John Platig, and Camila Lopes-Ramos for providing feedback on early drafts of this manuscript. DLD is supported by P01HL132825, P01HL114501 and a grant from the Alpha 1 Foundation. JQ is supported by a grant (R35CA220523) from the US National Cancer Institute. KG is supported by a grant (K25HL133599) from the US National Heart, Lung, and Blood Institute.

## Author contributions

KG and ARS developed the method, performed the analyses, and drafted the manuscript. DLD and JQ provided feedback on the analysis results and manuscript text. All authors have read and approved the final version of the manuscript.

## Supplementary Materials and Methods

### S1. Limitations to using genome-wide TFBS predictions when constructing gene regulatory networks

Position-weight-matrices (PWMs) are used to conceptualize the DNA sequence pattern bound by a transcription factor (TF). The genomic locations (“motif locations”) that match the pattern contained in a PWM can be used to predict a transcription factor’s binding sites (TFBSs). Multiple methods have been developed that leverage chromatin accessibility and other genomic information (e.g. sequence conservation, proximity to an annotated transcription start site, etc.) alongside PWMs to improve motif scoring and thereby TFBS predictions. These methods are generally benchmarked by comparing the scores associated with each motif location against a “gold standard” derived from ChIP-seq data. At a technical level, this amounts to comparing two vectors of length *Nm,* where *Nm* is the number of motif locations in the genome: (1) one containing scores associated with each motif location, and (2) one indicating whether or not there is also a ChIP-seq peak at these locations. This approach has two well-known limitations. First, by definition, this approach excludes locations in the genome where the TF is bound based on ChIP-seq data but there is no corresponding pattern in the DNA that matches the PWM. Secondly, because only a handful of motif locations are occupied by a transcription factor, these evaluations are prone to a high number of true-negatives, which can artificially inflate metrics such as the Area Under the Receiver-Operator Characteristic Curve (AUC-ROC, or more simply AUC).

These issues, especially the former, are exacerbated when constructing gene regulatory networks. In constructing gene regulatory networks, the motif locations that occur within the regulatory region (most often the promoter region) of a gene are often used to estimate which TFs regulate that gene. In contrast to genome-wide TFBS prediction, benchmarking the targets of a TF in a gene regulatory network involves comparing two vectors of length *N_g_*, where *N_g_* is the number of genes in the network: (1) one containing information regarding whether or not the TF’s motif was found within the regulatory region(s) of the genes, and (2) one containing information regarding whether or not there is a ChIP-seq peak for that same TF within the regulatory region(s) of the genes. Note that in this evaluation, genes whose regulatory regions contain a bound TF based on ChIP-seq data, but do not contain a corresponding match to that TF’s PWM, are no longer excluded, but instead are considered false negatives.

This shift has profound implications for model accuracy. Namely, even highly accurate TFBS predictions can, when naively used to construct a gene regulatory network as described above, lead to very poor network accuracy. To illustrate this seemingly counter-intuitive statement, **Supplemental Figure 1** depicts a toy example illustrating how the same data evaluated using either a ‘motif-centric’ or a ‘gene-centric’ approach can lead to dramatically different AUC values. The top panel shows three tracks that represent the genomic locations of (1) motifs, (2) gene promoters, and (3) open chromatin (for example, DNase-seq peaks). The middle panel shows the location of TF binding sites based on ChIP-seq peaks. In constructing this example, we selected parameters consistent with those observed in real biological data. Namely, 75% of gene promoters also contained an open chromatin region and 40% contained a predicted TFBS, 5% of predicted TFBS not in promoters were in regions of open chromatin (representing enhancers), and 25% of open chromatin regions contained a true TF binding site (ChIP-seq peak); no ChIP-seq peaks were located outside of open chromatin regions.

We computed the AUC using both a motif-centric and gene-centric approach. For the motif-centric comparison, motif locations that overlapped with open chromatin were given a score of “1” while motif locations that were not in open chromatin were given a score of “0”. Benchmarking this vector of motif scores against the ChIP-seq peaks resulted in a very high AUC value (AUC = 0.96; **Supplemental Figure 1**, first row of bottom panel). For the gene-centric approach we considered two scenarios. In the first, gene promoters that contained a motif were give a score of “1” and promoters that did not contain a motif were given a score of “0”. This is equivalent to using motif locations without chromatin information to predict a TF’s target genes, as in a regulatory network. In this set-up, the AUC value dropped severely (AUC = 0.55; **Supplemental Figure 1**, second row of bottom panel). For the second scenario, gene promoters that contained a motif that was also in open chromatin were given a score of “1” and all other gene promoters were given a score of “0”. In this case, despite the additional epigenetic information, there was only a minimal improvement in AUC (AUC = 0.63; **Supplemental Figure 1**, third row of bottom panel), and the accuracy was significantly below that obtained using the same information, but in a motif-centric manner.

This analysis shows that highly accurate results from motif-centric prediction approaches, including algorithms that combine PWM and chromatin accessibility information to predict TFBS, do not directly translate into accurate gene regulatory network models. This is not only due to a high number of true negatives, but also a high number of false negatives.

### S2. Details of the SPIDER method, input, and validation

#### S2.1 Details of the SPIDER algorithm

##### Step 1 – Intersect open chromatin regions with motif locations

In this step SPIDER uses bedtools^1^ (version v2.25.0) to intersect a BED file containing regions of open chromatin with a series of BED files containing the locations of TF motifs (one BED file per TF). The output of this step is a single BED file that contains the locations of TF motifs that are in open chromatin regions. By default, each of these locations is given a default score of one. Note that, in practice, the file produced by this step could be produced in another manner and still used by SPIDER.

##### Step 2 – Intersect motifs in open chromatin (from Step 1) with gene regulatory regions and create a seed regulatory network

Next, SPIDER uses bedtools to overlap a BED file containing the locations of TFs that are in open chromatin (created in Step 1) with a BED file containing the regulatory regions of genes. Note that a gene can have multiple associated regulatory regions in this second file. If a TF’s motif falls within the regulatory region(s) of a gene, then an edge is created between that TF and gene. The maximum score across all TF motif instances associated with a gene is used to weight the edge; by default, this value is one. The result of this step is an epigenetically-informed seed regulatory network between all transcription factors and genes.

##### Step 3 – Degree normalize seed network

The seed network from Step 2 consists of edges between *N_T_* TFs and *N_G_* genes. Let us denote the TF by gene adjacency matrix describing this network as *A*. Based on this matrix, we can calculate the average degree for each TF *i* as 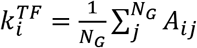, and the average degree for each gene *j* as 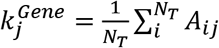. We use this information to degree normalize the seed network based on the transformation 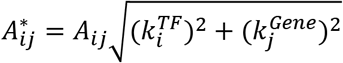.

##### Step 4 – Apply message passing

Finally, SPIDER applies the PANDA^2^ message-passing algorithm to the degree-normalized seed network *A\** calculated in Step 3. PANDA’s message-passing framework integrates information from three networks, representing TF protein-protein interaction *(P),* TF/gene regulation (*W* and gene co-expression (*C*). In SPIDER, *P* and *C* are set equal to the identity matrix; *W* is set equal to *A\**. PANDA returns a complete, bi-partite network with edge weights representing the likelihood that a TF regulates a gene; the distribution of these edge weights is similar to z-scores.

#### S2.2 Data used in SPIDER

The input data used by SPIDER includes BED files which contain (1) the genomic locations of potential transcription factor binding sites (one BED file per TF), (2) epigenetic (chromatin-accessibility) information, and (3) regulatory regions. In this study, we construct GRNs between 687 TFs and 27090 genes for six cell lines. We used hg19 coordinates for all the data in this study.

Identification of potential TFBS: In this project, we used motifs for 687 human transcription factors from the Catalog of Inferred Sequence Binding Preferences (Cis-BP)^3^ (http://cisbp.ccbr.utoronto.ca, accessed: July 7, 2015). We mapped the motifs to the human genome (hg19) using FIMO^4^, and retained all locations meeting a significance of p<10^-4^.

Epigenetic data: We obtained DNase-seq peak locations from ENCODE for 6 cell lines. Additional information about the DNase-seq data used, including lab and download URL, is given in **Supplemental File 1**.

Regulatory regions: We used RefSeq gene annotations downloaded from the UCSC genome browser (https://genome.ucsc.edu/cgi-bin/hgTables; accessed on 29^th^ May 2018) to define the regulatory regions of genes. We defined ranges of regulatory regions in terms of distance from the transcription start site (TSS): the proximal range was defined as window of 2kb centered around the TSS. For the analysis where we evaluated the potential of SPIDER to be used to predict distal regulatory events, we used pair of ranges, each with a width of 5kb. We chose separate distal windows of 5kb on both sides of the TSS in order to exclude promoter effects and to keep the density of regulatory seed network information similar between the promoter and distal analyses. For our primary distal analysis, these ranges were situated at a distance of 20-25kb both upstream and downstream of the gene transcriptional start site (TSS). We also evaluated SPIDER at various distance-ranges in the GM12878 and A549 cell lines. This included windows at ±5-10kb, ±20-25kb, ±45-50kb, ±70-75kb, and ±95-100kb around the TSS. We saw little variation in the accuracy of SPIDER predictions across these ranges.

#### S2.3 Validating SPIDER

ChIP-seq data for human transcription factors in six cell lines was obtained from ENCODE in narrow peak (BED) format. Information about the ChIP-seq data used, such as treatment, antibody, data freeze date, lab, and download URL, is provided in **Supplemental File 1**. For some TFs, multiple ChIP-seq experiments performed in the same cell line were available. In these cases, we made one composite file containing all ChIP-seq peaks using the bedtools merge function. To create gold standard networks from these data, ChIP-seq peaks were intersected with gene regulatory regions following the same protocol as used by SPIDER, described in Step 2 in **Section S2.1** above. We computed the AUC and AUPR of SPIDER predictions using the perfcurve function in matlab (R2014b). For the differential network analysis, the ‘negclass’ parameter for three class prediction problem was used.

### S3. Benchmarking SPIDER against other methods

We compared the performance of SPIDER to that of several other methods and available networks that incorporate epigenetic information. These fall into three main categories: (1) transcription factor binding site prediction methods (whose output needs to be processed to create gene regulatory networks comparable to those predicted by SPIDER); (2) publicly available gene regulatory networks (provided as a resource in other publications), and (3) gene regulatory network prediction methods. In analyzing these diverse resources, we processed the provided data in as consistent of manner as possible to support a fair comparison of the sources.

Below we describe each of these additional methods/networks and how we adapted their usage or output to systematically compare their performance with each other and with that of SPIDER. For all methods/networks we only focused on the subset of cell lines and transcription factors for which we had ChIP-seq data (see **Section S2.3** and **Supplemental File 1**). For each of the methods/networks we performed a threshold analysis to evaluate the accuracy of that method/network’s predictions. The thresholds used in these analyses were used to calculate the true positive rate, true negative rate, false positive rate, and false negative rate for each test performed, which is summarized in **Figure 5b** in the main text. The results reviewed in this section are summarized in **Supplemental Table 4** and illustrated in **Figure 5a** and **Supplemental Figure 4**.

#### Group 1: TFBS inference methods

We chose CENTIPEDE^5^ and PIQ^6^ as two representative, highly-cited examples of the many methods that have been developed to perform genome-wide TFBS prediction. Both of these were pioneering methods and are widely used as benchmarks.

##### CENTIPEDE

The CENTIPEDE algorithm identifies regions of the genome that are bound by transcription factors using a hierarchical Bayesian mixture model to integrate histone modifications and DNase I cleavage patterns with gene annotations and evolutionary conservation. We downloaded the data on TFBS predicted by CENTIPEDE from two resources:

a. the original CENTIPEDE repository: http://centipede.uchicago.edu/SimpleMulti/
b. website hosting supplemental material for^7^: https://noble.gs.washington.edu/proj/priors4search/

From the original CENTIPEDE repository, we downloaded the TFBS predictions for five cell lines (GM12878, H1HESC, HELAS3, HEPG2, K562) since these were analyzed in our primary SPIDER analysis. These data listed the genomic locations of predicted TFBS in hg18 coordinates, which we converted to hg19 coordinates using the liftOver program (downloaded from https://genome.ucsc.edu/cgi-bin/hgLiftOver, accessed October 2, 2018), but no scoring information. Therefore, to provide additional insight into the performance of CENTIPEDE, we also downloaded information regarding the posterior probability of TF binding data for the CENTIPEDE model, which was provided as part of the supplemental material in^7^. These data included six TFs in one exemplar cell line: GM12878. Although limited, these data included scores for all motif locations, allowing us to recapitulate the performance of CENTIPEDE at a genome-wide level and to gain insights into why the apparent superior performance of the algorithm did not directly translate into accurate network predictions (see **Section S1**).

To evaluate CENTIPEDE, we first focused on the limited dataset from^7^ and calculated the accuracy of the predicted motif locations based on PWM scores as well as the CENTIPEDE-predicted posterior probability scores. In particular, for each TF, we intersected the associated motif locations with the locations of ChIP-seq peaks (using the same ChIP-seq data as we used to benchmark SPIDER, see **Section S2.3**); locations that intersected with a ChIP-seq peak were considered true binding events. When then calculated the AUC based on the initial PWM scores associated with the motif locations as well as using the posterior probability scores predicted by CENTIPEDE. This analysis recapitulated CENTIPEDE’s excellent predictive performance, with a mean AUC of 0.9782 across the six TFs based on the posterior probability score. This compared to a mean AUC of only 0.7039 when using the original PWM scores (**Supplemental Figure 4a**).

Next, for each TF, we intersected motif locations with either proximal (±1kb around the TSS) or distal (±20-25kb around the TSS) gene regulatory regions (see Step 2 in **Section S2.1** above). Genes for which no associated regulatory region included a motif location were given a default weight of 0.001 less than the minimum posterior probability score across all motif locations, genes for which exactly one motif location was found across all associated regions were assigned the posterior probability score associated with that motif location, and genes for which more than one motif location was identified across all associated regulatory regions were assigned the highest of the posterior probability scores associated with the identified motif locations. For each TF, we compared the resulting vector of gene scores with information regarding whether or not there is also a ChIP-seq peak for that TF within at least one of the gene’s associated regulatory regions (see **Section S2.3**); we then used this information to calculate both an AUC and AUPR for that TF. We found that the mean AUC across the six TFs was only 0.5403 for the proximal regions (**Supplemental Figure 4b-c**) and 0.5632 for the distal regions (**Supplemental Figure e-f**).

We also performed a threshold analysis since this would better align with the data supplied in the original CENTIPEDE repository. In particular, we identified motif locations that had a high posterior probability (score > 5) and intersected those locations with proximal and distal gene regulatory regions (as in Step 2 of SPIDER, see **Section S2.1**). Genes which were not associated with any regulatory regions that included one of these high-scoring motif locations were given a default weight of zero, while genes associated with at least one regulatory region that contained one or more of these high-scoring motif locations were given a score of one. The mean AUC across the six TFs in this context was only 0.5236 for the proximal regions and 0.5189 for the distal regions (**Supplemental Table 4**).

Finally, we analyzed the data provided from original CENTIPEDE repository, which contained bed files listing only high-probability motif locations. Exactly as in the above analysis, we intersected motif locations with gene regulatory regions, and then benchmarked each resulting profile with its corresponding standard based on ChIP-seq data for that TF and cell-line (47 total comparisons). We observed overall poor predictive performance, with a mean AUC of only 0.5156 for proximal regions and 0.5218 for distal regions. These values are very similar to those we obtained when analyzing the more detailed data from only six TFs, as described above. The results of these analyses are shown in **Figure 5a** (proximal) and **Supplemental Figure 4d** (distal).

##### Protein-DNA Interaction quantitation (PIQ)

PIQ is a tool for global TF binding site detection using a combination of motif information and DNase-seq data^6^. Similar to CENTIPEDE, PIQ estimates binding probabilities associated with a set of motif locations. We downloaded data containing PIQ predictions from http://piq.csail.mit.edu/data/141105-3618f89-hg19k562.calls/. These data contained predictions for the K562 cell line and included 48 motifs that were also associated with one of the TFs for which we had K562 ChIP-seq data; these 48 motifs corresponding to 25 unique TFs.

As in our CENTIPEDE analysis (see above), we first evaluated the overall accuracy of the PIQ scores across all motif locations. More specifically, for each motif, we merged together the genomic locations provided for both the standard and reverse compliment PIQ analyses. We then intersected these combined locations with the locations of ChIP-seq peaks (using the same ChIP-seq data as we used to benchmark SPIDER, see **Section S2.3**); locations that intersected with a ChIP-seq peak were considered true binding events. We used these data to calculate an AUC based on the initial PWM scores as well as based the scores assigned by PIQ. In this context, PIQ performed well, with a mean AUC of 0.7729 across the 48 motifs based on the PIQ score, compared to a mean AUC of 0.6001 for the PWM score (**Supplemental Figure 4a**).

Next, we intersected motif locations with either proximal or distal regulatory regions (as described in Step 2 of SPIDER, see **Section S2.1** above). Exactly as in our CENTIPEDE analysis (see above), genes whose regulatory regions didn’t include a motif location were given a default weight of 0.001 less than the minimum score across all motif locations, genes associated with exactly one motif location across all associated regulatory regions were assigned the score associated with that motif location, and genes with more than one motif location identified across all associated regulatory regions were assigned the highest score across all associated motif locations. For each motif, we compared the resulting vector of gene scores with information regarding whether or not there is also a ChIP-seq peak for the corresponding TF within at least one of the gene’s associated regulatory regions; this allowed us to calculate an AUC and AUPR for that motif. We found that the mean AUC across the 48 motifs was only 0.5703 for the proximal regions (**Supplemental Figure 4b-c**) and 0.5606 for the distal regions (**Supplemental Figure e-f**).

Finally, we also performed a threshold analysis where we identified motif locations that had a high score (score > 0) and intersected those locations with proximal and distal gene regulatory regions (as in Step 2 of SPIDER, see **Section S2.1**). Genes whose associated regulatory regions didn’t include a high-scoring motif location were given a default value of zero, while genes that were associated with at least one regulatory region that contained one or more high-scoring motif locations were given a score of one. The mean AUC across the 48 tested motifs was 0.5559 for proximal regions and 0.5389 for distal regions. The results of these analyses are shown in **Figure 5a** (proximal) and **Supplemental Figure 4d** (distal).

Overall, these analyses indicate that the predictions from CENTIPEDE and PIQ cannot be naively used for regulatory network inference. TFBS prediction approaches that combine motif locations with chromatin data are poor predictors of TF occupancy when restricted to gene regulatory regions. For more information, see **Section S1**, above.

#### Group 2: Available gene regulatory networks

##### Networks published along with Neph et. al.^8^

One of the first attempts to use DNase footprints to construct regulatory networks was published by Neph et. al.^8^. In this paper, the authors combined TF motif locations with genome-wide DNase1 footprints to construct regulatory networks of transcription factors. Specifically, if the DNase1 footprint of transcription factor *i* was identified within the promoter region of the gene encoding transcription factor *j,* a regulatory edge was drawn from transcription factor *i* to transcription factor *j.* By design, only genes that encode transcription factors were included in these networks.

We downloaded the regulatory networks associated with^8^ from http://www.regulatorynetworks.org. From these, we selected the networks for the K562 and HEPG2 cell lines, since those were included in the six cell lines for which we had ChIP-seq data. Note that the downloaded networks were binary (non-weighted), only contained transcription factors, and were restricted to promoter-based regulatory predictions. We evaluated the accuracy of each of the TFs within these networks using the ChIP-seq gold standards used to evaluate TFs in the proximal SPIDER networks (see **Section S2.3**), restricting those standards to only include target genes within in the Neph et. al. networks (i.e. the subset of genes encoding transcription factors). The mean AUC across all the tests performed was 0.5178, similar to mean AUC for threshold proximal CENTIPEDE analysis. The results of this analysis are illustrated in **Figure 5a** in the main text.

##### Networks published along with Marbach et. al.^9^

Another collection of gene regulatory network models was provided along with the article published by Marbach et. al.^9^. These network models were constructed by identifying the promoters and enhancers of genes using CAGE expression profiles, and then linking TF motif locations found within these regions to the corresponding target genes.

We downloaded the networks constructed by Marbach et. al.^9^ from http://regulatorycircuits.org. Among the 394 networks provided through this resource, we identified five for which we had corresponding ChIP-seq data and could use for benchmarking and validation. The downloaded networks were binary (non-weighted), limited to the set of genes used for network building in the CAGE analysis (these genes were provided by the authors in a supplemental file together with the downloaded networks), and did not distinguish between proximal and distal regulation. Similar to how we evaluated the networks associated with Neph et. al. (see above), we evaluated the accuracy of these networks using the ChIP-seq gold standards used to evaluate the proximal SPIDER networks (see **Section S2.3**). Specifically, we restricted each gold standard to only include the set of genes used in the CAGE analysis and evaluated each TF in each network using its corresponding TF and cell line ChIP-seq standard (167 total tests). The mean AUC across all the tests performed was 0.5595, similar to the mean AUC for the threshold proximal PIQ analysis. The results of this analysis are illustrated in **Figure 5a** in the main text.

#### Group 3: Gene regulatory network prediction methods

##### TEPIC

TEPIC performs a segmentation-based method to predict TF-binding by the combining PWMs and openchromatin assay data. TEPIC computes TF binding affinities based on a biophysical model of TF binding which detects low affinity binding sites. Of all the methods/networks evaluated, the TEPIC input information and approach is the most similar to that of SPIDER.

##### Reconstructing networks using TEPIC

We downloaded the TEPIC program and supporting files from https://github.com/SchulzLab/TEPIC. To create regulatory predictions, the input to TEPIC includes (1) a set of formatted PWMs, (2) the full genomic sequence in fasta format, (3) open chromatin regions in BED format, (4) a gene annotation file that indicates the location of gene transcriptional start sites, and (5) a window size to indicate the range around the TSS to designate as the regulatory region. For (1) we used a set of formatted JASPAR motifs (“human_jaspar_hoc_kellis.PSEM”) provided with the TEPIC program. For (2) we used the same hg19 fasta sequence file used for the FIMO motif scan we performed to create the SPIDER input files (see **Section S2.2** above, “Identification of potential TFBS”). For (3) we used the same DNase peak bed files we used when running SPIDER (see **Section S2.2** above, “Epigenetic Data”).

For (4) we created custom gene annotation files that would ensure TEPIC used the same regulatory regions that we used when running SPIDER. In particular, for the proximal regions we created a gtf file with each entry corresponding to a unique TSS. We note that some genes are associated with more than one TSS; however, TEPIC by default only considers one entry for each unique gene symbol. Therefore, in order for TEPIC to treat these as separate regulatory regions (as we did in SPIDER), we listed a gene (“GeneName”) that is annotated to multiple TSS as GeneName::COPY1, GeneName::COPY2, etc, one copy per unique TSS. When running TEPIC for proximal regions we set the window size to 1000, which corresponded to running TEPIC on regulatory regions defined as ±1kb around each TSS – the same regions we used for our proximal SPIDER analysis.

To run TEPIC using distal regions, we created a gtf file that had two entries for each unique TSS, one noting a location 22500bp upstream of the TSS and one noting a location 22500bp downstream of the TSS. These locations correspond to the mid-points of the regulatory regions we used when performing our distal analysis with SPIDER. In order for TEPIC to treat these as separate regulatory regions, each gene (“GeneName”) was listed twice using the convention GeneName::U and GeneName::D, and annotated to the upstream and downstream TSS locations, respectively. As in the proximal analysis, genes that were associated with more than one TSS were denoted by GeneName::U::COPY1, GeneName::U::COPY2, etc. When running TEPIC for distal regions we set the window size to 2500, which corresponded to running TEPIC on regulatory regions located at ±20-25kb around each TSS – the same regions we used for our distal SPIDER analysis.

We ran TEPIC using DNase data for each of the six cell lines and gene annotations for proximal as well as distal regulatory regions (twelve total analyses). To handle multiple regulatory regions associated with the same gene (eg a gene associated with multiple associated TSS), for each TF motif to gene edge, we weighted the edge based on its maximum score across all possible inferences. Note that only genes associated with at least one regulatory region that overlaps with a DNase peak are contained in the TEPIC output file. Therefore, for completeness, edges targeting genes not in the TEPIC output were given a default score of zero; this ensured the networks we inferred using TEPIC contained the same set of target genes as the networks inferred using SPIDER.

##### Analyzing TEPIC-predicted networks

We first evaluated the accuracy of TEPIC predictions using continuous scores. Specifically, for each TF motif in each cell line, we extracted the vector of edge-weights from that motif to all genes in the TEPIC-predicted network and compared that vector to the TF’s targets in the corresponding cell line specific ChIP-seq gold standard (see **Section S2.3**). Based on this analysis, we observed a mean AUC across all the tests of 0.7405 for the proximal analysis (**Supplemental Figure 4b-c**) and 0.7586 for the distal analysis (**Supplemental Figure 4e-f**). Overall the predictions made by TEPIC were more accurate than all the other methods/networks we evaluated, with the exception of SPIDER

We also performed a threshold analysis. For each cell line network, we selected the top 25% of edges by weight. We evaluated the accuracy of each TF motif in these binarized networks using the corresponding TF and cell line specific ChIP-seq gold standard. Thresholding the TEPIC results negatively impacted its performance. When thresholding, we observe a mean AUC of only 0.5820 across all tests for the proximal analysis and a mean AUC of 0.6549 for the distal analysis. The results of this threshold analysis are shown in **Figure 5a** of the main text.

##### SPIDER

Our main evaluation of SPIDER’s performance involved benchmarking the continuous edge scores predicted by SPIDER against ChIP-seq predictions. The results of these evaluations for individual TFs is shown in **Figures 2 & 4** (proximal and distal AUC, respectively) in the main text and in **Supplemental Figures 2 & 3** (proximal and distal AUPR, respectively). These results are summarized across all cell lines and shown in **Supplemental Figure 4b-c** (proximal networks) and **Supplemental Figure 4e-f** (distal networks) alongside the subset of methods for which we could also perform continuous evaluations (six motifs in CENTIPEDE, PIQ, and TEPIC). For SPIDER, we observe a mean AUC of 0.7949 (proximal) and 0.8477 (distal) across all tests.

However, for many of the methods/networks described above we were only able to evaluate binary predictions. Therefore, to support a more equitable comparison of SPIDER to these other methods/networks, we also performed a threshold analysis based on SPIDER’s predictions. As in the TEPIC evaluations (see above), for each cell line network, we identified the top 25% of edges by weight. We evaluated the accuracy of each TF in these binarized networks using the corresponding TF and cell line specific ChIP-seq gold standard. We observed a mean AUC of only 0.6950 across all tests for the proximal analysis and a mean AUC of 0.7747 for the distal analysis. Although these AUC values are lower than the ones we obtained when evaluating SPIDER’s continuous scores, they are still much higher than those we obtained from all other sources in the context of thresholding. The results of this threshold analysis are shown in **Figure 5a** of the main text.

##### SPIDER input data

For our final evaluation, we also determined the accuracy of the FIMO motif scan (see **Section S2.2** “Data used in SPIDER”). This is an important evaluation since the quality of the data used to seed SPIDER may impact the algorithm’s performance and explain its higher accuracy compared to the other sources we evaluated. To begin, we intersected the genomic locations of each motif with each of the ChIP-seq bed files corresponding to that motif’s associated TF. Locations with an overlapping ChIP-seq peak were considered true binding events. Separately, we also intersected the genomic locations of each motif with open chromatin regions for each of the six cell lines, as defined in the DNase BED files used by SPIDER. Motif locations that overlapped with open chromatin regions were given a score of one, and all other locations were assigned a score of zero. This mirrors Step 1 of the SPIDER algorithm (see **Section S2.1**).

We used these data to calculate an AUC based on the initial PWM scores as well as based the binary “scores” obtained from overlapping with chromatin information. We found that, at a genome wide level, the data used as an input to SPIDER was already fairly accurate, with a mean AUC across all tests of 0.6596 based on PWM scores, and a mean AUC across all tests of 0.7431 based on the binary DNase-based scoring scheme. These results are illustrated in **Supplemental Figure 4a**. Overall, the accuracy of the motif scan that we used to construct the seed networks for SPIDER appears fairly similar to that used by CENTIPEDE and PIQ. Thus, the accuracy of the SPIDER input data cannot explain its highly superior performance compared to these other approaches in the gene regulatory network context.

### S4. Detecting novel interactions inferred by SPIDER

For each cell line network, we selected TF-gene relationships that were absent in the SPIDER seed network. Because SPIDER uses PANDA to perform message-passing, the weights of these edges can be interpreted as Z-scores^2^. Therefore, to identify significant edges in this class, we converted the weights of edges into probabilities using the *pnorm()* function in R and corrected for multiple comparisons using the Benjamini-Hochberg method. We selected top significant edges for further analysis using an FDR cutoff of 0.05. For the A549 proximal network this corresponded to an edge weight cutoff of 4.64.

## Supplemental Tables

**Supplemental Table 1:** Statistical evaluation of the increase in AUC for the SPIDER Regulatory Networks (SRN) compared to the naïve regulatory networks (NRN) and SPIDER seed networks (SSN). Density for the naïve seed (NSN) network was 39.73%.

| Proximal Region<br>(0 – 1kb from TSS) |  |  |  |  |  |  |  |  |  |
| --- | --- | --- | --- | --- | --- | --- | --- | --- | --- |
| Cell line<br>(SSN<br>density) | AUC values (95% CI) |  |  | SRN vs NRN |  |  | SRN vs SSN |  |  |
|  | NRN | SSN | SRN | Percent<br>increase | z-score<br>(DeLong's<br>test) | p-value | Percent<br>increase | z-score<br>(DeLong's<br>test) | p-value<br>(two-<br>sided) |
| **A549**<br>**(5.89%)** | **0.574**<br>(0.5722-0.5763) | **0.594**<br>(0.5921-0.595) | **0.816**<br>(0.8144-0.817) | 42.1 | -220.78 | < 2.2e-16 | 37.3 | -316.34 | < 2.2e-16 |
| **H1-HESC**<br>**(6.04%)** | **0.597**<br>(0.5957-0.5991) | **0.607**<br>(0.6057-0.6083) | **0.726**<br>(0.7248-0.7277) | 21.6 | -138.43 | < 2.2e-16 | 19.6 | -194.41 | < 2.2e-16 |
| **HELAS3**<br>**(5.64%)** | **0.579**<br>(0.5725-0.5755) | **0.602**<br>(0.6008-0.603) | **0.8093**<br>(0.8072-0.8092) | 39.7 | -289.91 | < 2.2e-16 | 34.3 | -412.16 | < 2.2e-16 |
| **HEPG2**<br>**(5.49%)** | **0.583**<br>(0.5778-0.5806) | **0.598**<br>(0.5974-0.5994) | **0.770**<br>(0.77-0.7721) | 32.1 | -248.52 | < 2.2e-16 | 28.7 | -344.19 | < 2.2e-16 |
| **GM12878**<br>**(5.77%)** | **0.581**<br>(0.5771-0.5795) | **0.602**<br>(0.6014-0.6031) | **0.819**<br>(0.8178-0.8193) | 40.9 | -373.18 | < 2.2e-16 | 36.0 | -538.96 | < 2.2e-16 |
| **K562**<br>**(5.69%)** | **0.578**<br>(0.5738-0.5761) | **0.596**<br>(0.5956-0.5972) | **0.795**<br>(0.7943-0.7959) | 37.5 | -358.64 | < 2.2e-16 | 33.3 | -505.68 | < 2.2e-16 |

**Supplemental Table 2:** Characteristics of the input data as well as output predictions made for distal ranges.

| Ranges | GM12878 |  |  | A549 |  |  |
| --- | --- | --- | --- | --- | --- | --- |
|  | AUC - SRN | AUC- SSN | Density (%) | AUC - SRN | AUC- SSN | Density (%) |
| **±0-1kb** | 0.819 | 0.602 | 5.77 | 0.815 | 0.593 | 5.89 |
| **±5-10kb** | 0.842 | 0.622 | 5.89 | 0.846 | 0.629 | 4.84 |
| **±20-25kb** | 0.855 | 0.620 | 4.97 | 0.849 | 0.627 | 4.73 |
| **±45-50kb** | 0.854 | 0.620 | 5.12 | 0.848 | 0.627 | 4.82 |
| **±70-75kb** | 0.853 | 0.620 | 5.10 | 0.846 | 0.624 | 4.77 |
| **±95-100kb** | 0.849 | 0.618 | 5.04 | 0.849 | 0.625 | 4.68 |

**Supplemental Table 3:**
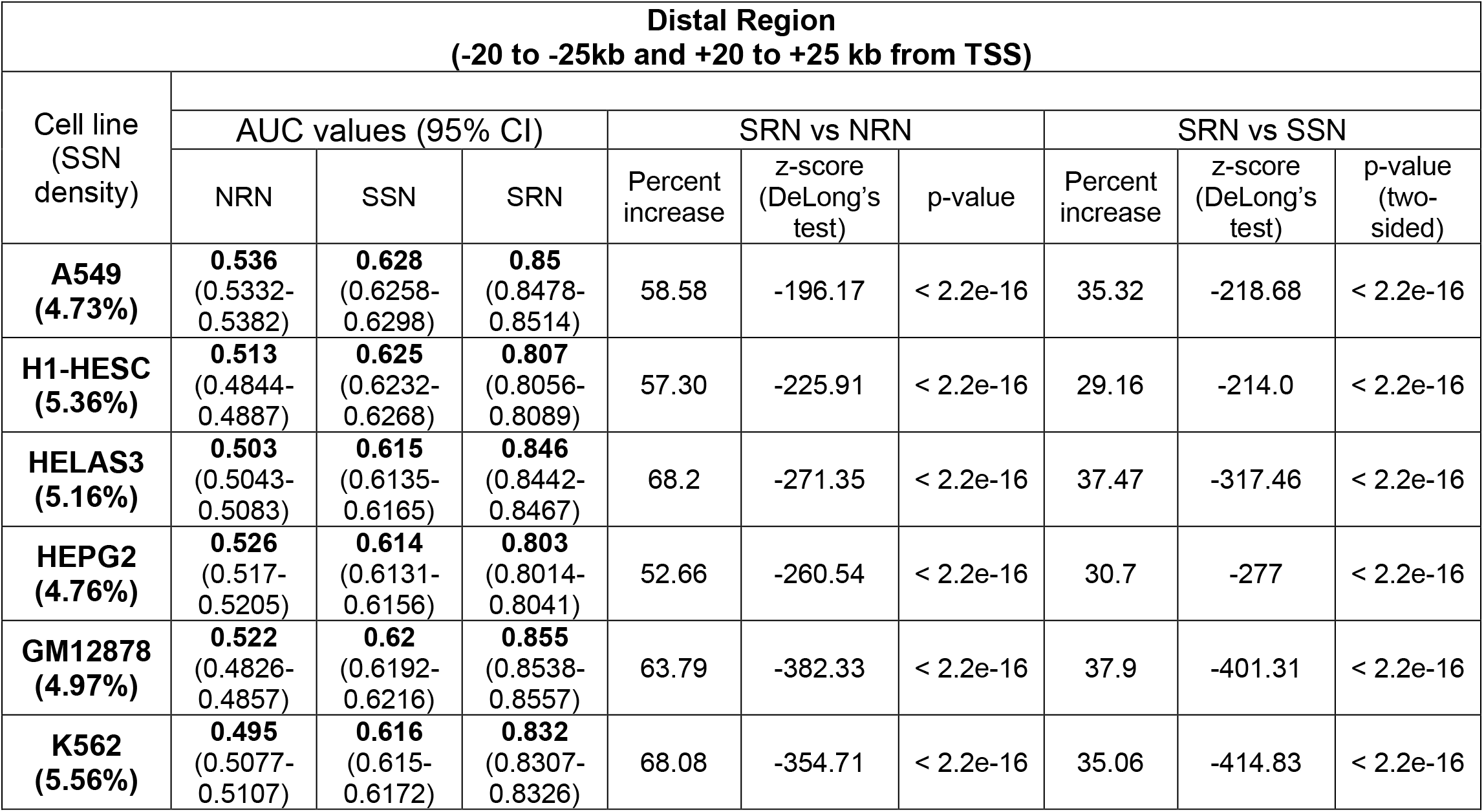
Statistical evaluation of the increase in AUC for the SPIDER Regulatory Networks (SRN) compared to the naïve regulatory networks (NRN) and SPIDER seed networks (SSN), based on a distal range of ±20-25kb around the TSS. Density for the naïve seed network (NSN) was 80.52%.

**Supplemental Table 4:** Results when evaluating predicted TF–gene relationships estimated or provided by various sources. The number of tests performed for each source are listed as well as the mean AUC and AUPR values across all tests. Note that not all evaluations could be performed in all sources. In particular, continuous AUC and AUPR could not be calculated for many sources. CENTIPEDE-sub refers to a small subset of CENTIPEDE predictions for which we had continuous prediction information. CL=cell line; TF=transcription factor; thresh = threshold analysis; cont = continuous analysis; PWM = position weight matrix score used; * = sources shown in **Figure 5**;^†^ = sources shown in **Supplemental Figure 4d**.

|  | Characteristics |  |  | Average AUC Across Tests |  |  |  |  |  | Average AUPR Across Tests |  |  |  |
| --- | --- | --- | --- | --- | --- | --- | --- | --- | --- | --- | --- | --- | --- |
|  | # Evaluated |  |  | proximal |  | distal |  | genome-wide |  | proximal | distal | genome-wide |  |
| Sources | TFs | CLs | Tests | thresh | cont | thresh | cont | PWM | Score | cont | cont | PWM | Score |
| CENTIPEDE-sub | 6 | 1 | 6 | 0.524 | 0.540 | 0.519 | 0.563 | 0.704 | 0.978 | 0.316 | 0.199 | 0.303 | 0.730 |
| CENTIPEDE*† | 19 | 5 | 47 | 0.516 | X | 0.522 | X | X | X | X | X | X | X |
| PIQ*† | 25 | 1 | 48 | 0.556 | 0.570 | 0.539 | 0.561 | 0.600 | 0.773 | 0.280 | 0.141 | 0.066 | 0.229 |
| Neph* | 64 | 2 | 91 | 0.518 | X | X | X | X | X | X | X | X | X |
| Marbach* | 74 | 5 | 167 | 0.559 | X | X | X | X | X | X | X | X | X |
| TEPIC*† | 97 | 6 | 243 | 0.582 | 0.741 | 0.655 | 0.759 | X | X | 0.284 | 0.221 | X | X |
| SPIDER*† | 103 | 6 | 254 | 0.695 | 0.795 | 0.775 | 0.848 | X | X | 0.374 | 0.332 | X | X |
| SPIDER Input | 103 | 6 | 254 | 0.601 | X | 0.622 | X | 0.660 | 0.743 | X | X | 0.072 | 0.182 |
| Figure # | X | X | X | 5 | S4b | S4d | S4e | S4a | S4a | S4c | S4f | X | X |

**Supplemental Table 5:** The frequency of top genes in significant SPIDER edges (FDR<0.05) that were not part of the SPIDER seed network, for each of the six cell lines and for the proximal and distal regions.

| Cell line | Most frequent genes among novel edges inferred (FDR p<0.05) by SPIDER |  |  |  |
| --- | --- | --- | --- | --- |
| | Proximal region ($\pm 0$-1kb) | | Distal region ($\pm 20$-25kb) | |
| A549 | PDE4D | 200 | MIR6870 | 378 |
|  | MIR100HG | 132 | GPR25 | 375 |
|  | ZBTB20 | 108 | CABLES1 | 369 |
|  | TGIF1 | 85 | NINJ2 | 369 |
|  | LINC-PINT | 31 | SEPT9 | 368 |
|  | NCOA7 | 29 | NFKBIZ | 366 |
|  | MIA2 | 11 | MIR100HG | 362 |
|  | ABLIM1 | 9 | SERTAD3 | 362 |
|  | FAM107B | 5 | MIR663B | 360 |
| H1-HESC | No significant edges |  | CXXC5 | 241 |
|  |  |  | HOXA3 | 211 |
|  |  |  | SEPT9 | 209 |
|  |  |  | BDNF | 134 |
|  |  |  | FGR | 116 |
| HELA-S3 | MIA2 | 163 | PPARG | 297 |
|  | ZBTB20 | 139 | PLEC | 286 |
|  | PLEC | 35 | CASC21 | 273 |
|  | MIR99AHG | 25 | KRT8 | 271 |
|  | UBE2D3 | 2 | SMAD3 | 267 |
|  | CXXC5 | 1 | PDE4D | 257 |
|  | DNAJB4 | 1 | CASC19 | 253 |
| HEPG2 | No significant edges |  | CFLAR | 288 |
|  |  |  | NDUFS2 | 271 |
|  |  |  | SYS1 | 268 |
|  |  |  | CXXC5 | 205 |
|  |  |  | PCBD2 | 203 |
|  |  |  | FAM50B | 199 |
|  |  |  | KIFC3 | 197 |
| GM12878 | SEPT9 | 106 | SLC44A2 | 376 |
|  | ZFP36L1 | 2 | LOC100120357 | 375 |
|  |  |  | TMEM151B | 375 |
|  |  |  | MXRA7 | 373 |
|  |  |  | SPTBN1 | 372 |
| K562 | FOXP1 | 84 | CXXC5 | 290 |
|  | MAPRE2 | 1 | MIR4761 | 281 |
|  | SAMSN1 | 1 | PHF19 | 277 |
|  |  |  | COMT | 268 |
|  |  |  | RNY4 | 262 |

**Supplemental Table 6:** For each of the six cell lines, the top edges (by weight) that were predicted by SPIDER around TFs for which we had ChIP-seq data in that cell line. Top edges that either had, or did not have, motif evidence are shown. Light blue shading indicates a true positive in the seed network that was retained after running SPIDER; light red shading indicates a false negative in the seed network that was recovered by SPIDER.

|  | Cell line | Range | TF | Gene | SRN | Motif | ChIP-seq |
| --- | --- | --- | --- | --- | --- | --- | --- |
| 1. | A549 | 0-1kb | MAX | MIR100HG | 22.35 | Yes | No |
|  |  |  | CEBPB | PDE4D | 21.07 | Yes | Yes |
|  |  |  | SIX5 | PDE4D | 20.22 | Yes | Yes |
|  |  |  | CREB1 | PDE4D | 10.03 | No | Yes |
|  |  |  | ETS1 | MIR100HG | 8.86 | No | No |
|  |  |  | MAX | PDE4D | 6.86 | No | Yes |
|  | A549 | 20- 25kb | MAX | PDE4D | 38.50 | Yes | Yes |
|  |  |  | ETS1 | PDE4D | 34.85 | Yes | Yes |
|  |  |  | CREB1 | PDE4D | 32.49 | Yes | Yes |
|  |  |  | YY1 | PDE4D | 14.61 | No | No |
|  |  |  | ETS1 | SMAD3 | 12.85 | No | Yes |
|  |  |  | CREB1 | ITGB1 | 12.25 | No | Yes |
| 2. | H1HESC | 0-1kb | SP1 | CNGA4 | 19.69 | Yes | No |
|  |  |  | SRF | FOXP1 | 18.86 | Yes | Yes |
|  |  |  | SP1 | SUGT1P1 | 18.77 | Yes | No |
|  |  |  | ZNF143 | PAX6 | 3.76 | No | Yes |
|  |  |  | SIX5 | PAX6 | 3.48 | No | No |
|  |  |  | BACH1 | PAX6 | 3.22 | No | Yes |
|  | H1-HESC | 20-25kb | SRF | CXXC5 | 25.12 | Yes | No |
|  |  |  | MAX | CXXC5 | 23.53 | Yes | No |
|  |  |  | SRF | HOXA3 | 22.12 | Yes | Yes |
|  |  |  | TEAD4 | CXXC5 | 8.44 | No | Yes |
|  |  |  | CEBPB | CXXC5 | 8.43 | No | No |
|  |  |  | NANOG | HOXA3 | 8.29 | No | No |
| 3. | HELA-S3 | 0-1kb | E2F4 | MIA2 | 24.76 | Yes | No |
|  |  |  | E2F1 | MIA2 | 21.28 | Yes | No |
|  |  |  | MAZ | TRANK1 | 19.75 | Yes | No |
|  |  |  | E2F1 | ZBTB20 | 6.96 | No | Yes |
|  |  |  | ZNF143 | MIA2 | 6.51 | No | No |
|  |  |  | TCF7L2 | MIA2 | 6.18 | No | No |
|  | HELA-S3 | 20-25kb | E2F4 | KRT8 | 29.19 | Yes | No |
|  |  |  | RFX5 | PDE4D | 26.52 | Yes | Yes |
|  |  |  | STAT1 | PDE4D | 26.19 | Yes | Yes |
|  |  |  | E2F1 | PDE4D | 13.12 | No | No |
|  |  |  | ZNF143 | PDE4D | 11.90 | No | No |
|  |  |  | MAX | PDE4D | 11.77 | No | Yes |
| 4. | HEPG2 | 0-1kb | MYBL2 | FOXP1 | 21.91 | Yes | No |
|  |  |  | MAZ | DKFZP434H168 | 20.12 | Yes | No |
|  |  |  | SRF | PDE4DIP | 19.78 | Yes | No |
|  |  |  | HSF1 | FOXP1 | 4.97 | No | No |
|  |  |  | HSF1 | MIA2 | 4.18 | No | No |
|  |  |  | HSF1 | MIR4461 | 4.17 | No | No |
|  | HEPG2 | 20-25kb | BRCA1 | CFLAR | 28.18 | Yes | No |
|  |  |  | SRF | CFLAR | 27.36 | Yes | Yes |
|  |  |  | HSF1 | CFLAR | 25.10 | Yes | No |
|  |  |  | MYBL2 | CFLAR | 10.32 | No | Yes |
|  |  |  | SRF | NDUFS2 | 8.02 | No | Yes |
|  |  |  | SRF | SYS1 | 7.97 | No | No |
| 5. | GM12878 | 0-1kb | MAX | SEPT9 | 24.35 | Yes | Yes |
|  |  |  | STAT5A | SEPT9 | 20.15 | Yes | Yes |
|  |  |  | EBF1 | SEPT9 | 20.07 | Yes | Yes |
|  |  |  | IKZF1 | SEPT9 | 7.6 | No | No |
|  |  |  | ETS1 | SEPT9 | 6.24 | No | Yes |
|  |  |  | ELK1 | SEPT9 | 5.55 | No | No |
|  |  | 20-25kb | ELK1 | CXXC5 | 30.49 | Yes | Yes |
|  |  |  | SRF | CXXC5 | 30.26 | Yes | No |
|  |  |  | STAT3 | CXXC5 | 28.86 | Yes | Yes |
|  |  |  | IKZF1 | CXXC5 | 15.07 | No | Yes |
|  |  |  | SIX5 | CXXC5 | 14.46 | No | No |
|  |  |  | PAX5 | CXXC5 | 14.15 | No | Yes |
| 6. | K562 | 0-1kb | SRF | FOXP1 | 22.86 | Yes | No |
|  |  |  | ZNF263 | SKAP1 | 19.19 | Yes | No |
|  |  |  | MAZ | DKFZP434H168 | 19.18 | Yes | Yes |
|  |  |  | ZNF143 | FOXP1 | 6.47 | No | No |
|  |  |  | ETS1 | FOXP1 | 6.22 | No | No |
|  |  |  | THAP1 | FOXP1 | 6.15 | No | No |
|  |  | 20-25kb | MAX | CXXC5 | 27.59 | Yes | Yes |
|  |  |  | ETS1 | COMT | 26.68 | Yes | Yes |
|  |  |  | ELK1 | CXXC5 | 26.34 | Yes | Yes |
|  |  |  | STAT5A | COMT | 10.52 | No | Yes |
|  |  |  | MAX | COMT | 10.41 | No | Yes |
|  |  |  | E2F4 | CXXC5 | 10.14 | No | Yes |

**Supplemental Figure 1:**
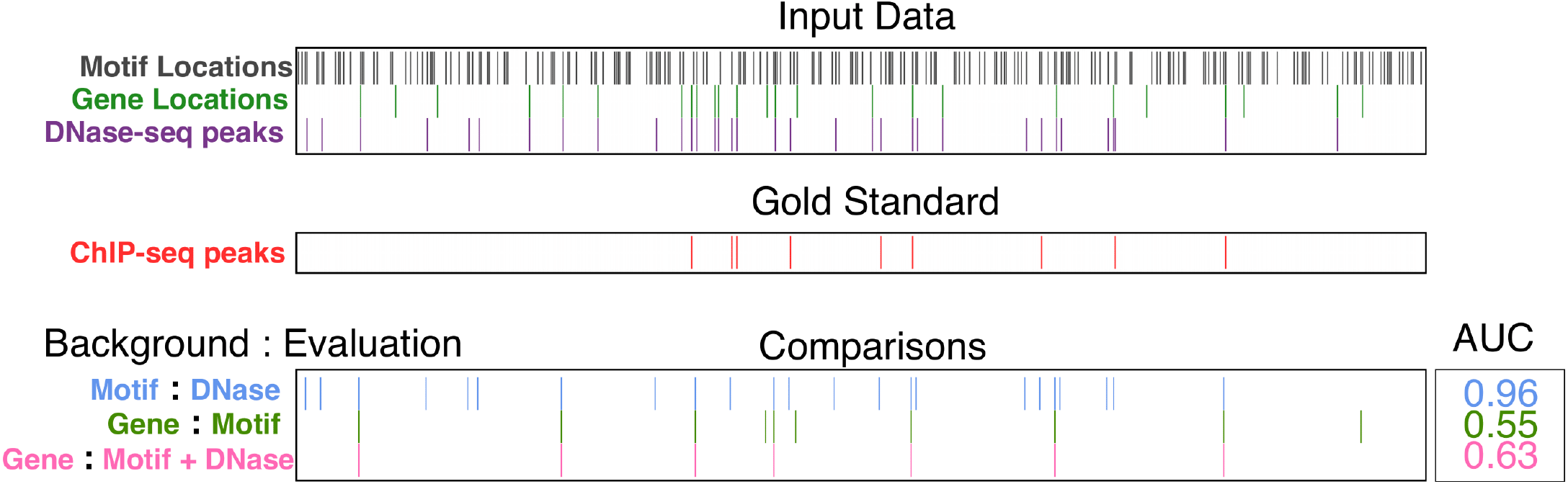
Illustrative example of how TFBS predictions are often used to estimate gene regulation, and the impact on prediction accuracy. The top panel shows the genomic locations of TF motifs, gene promoters, and open chromatin regions. The second panel shows the true locations of TF binding based on ChIP-seq data. The bottom panel illustrates combinations of data from the top panel. These combinations were evaluated using a benchmark derived from the ChIP-seq data in the middle panel. The word to the left of the colon indicates the set of elements being evaluated, while the word to the right indicates what is considered a positive prediction. For example, in th’e final row, each *gene* is assigned a value of 1 or 0 based on whether or not that gene’s promoter also contains both a *motif* and a *DNase-seq* peak; this is benchmarked against information regarding whether or not each gene’s promoter contains a true binding event, based on ChIP-seq peaks. In this toy example, we see that the AUC value in this evaluation is 0.63.

**Supplemental Figure 2:**
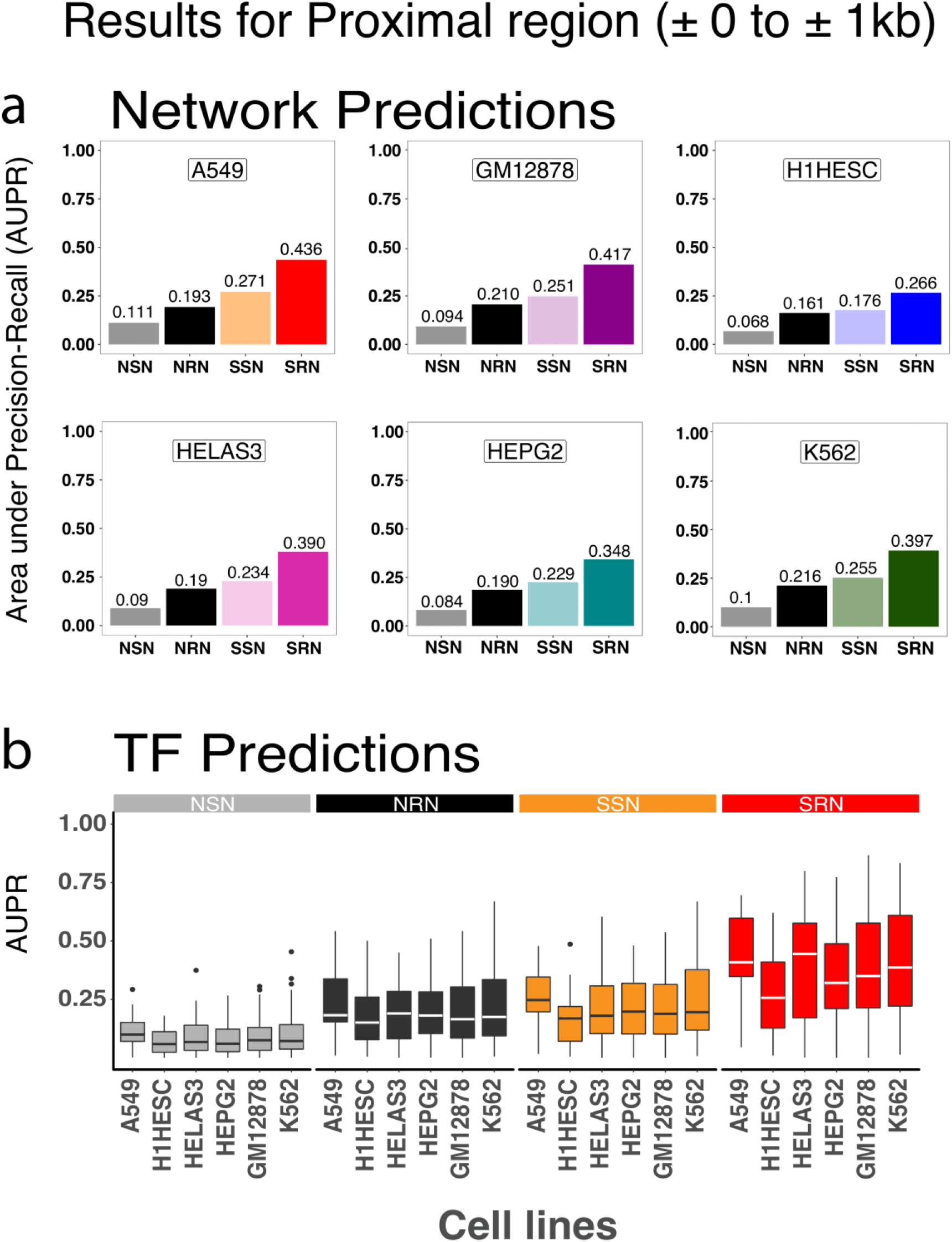
AUPR predictions for the proximal region. **a.** AUPR scores for four types of networks in six cell-lines. **b.** Distribution of AUPR scores across individual TFs in these networks. See also **Figure 2**.

**Supplemental Figure 3:**
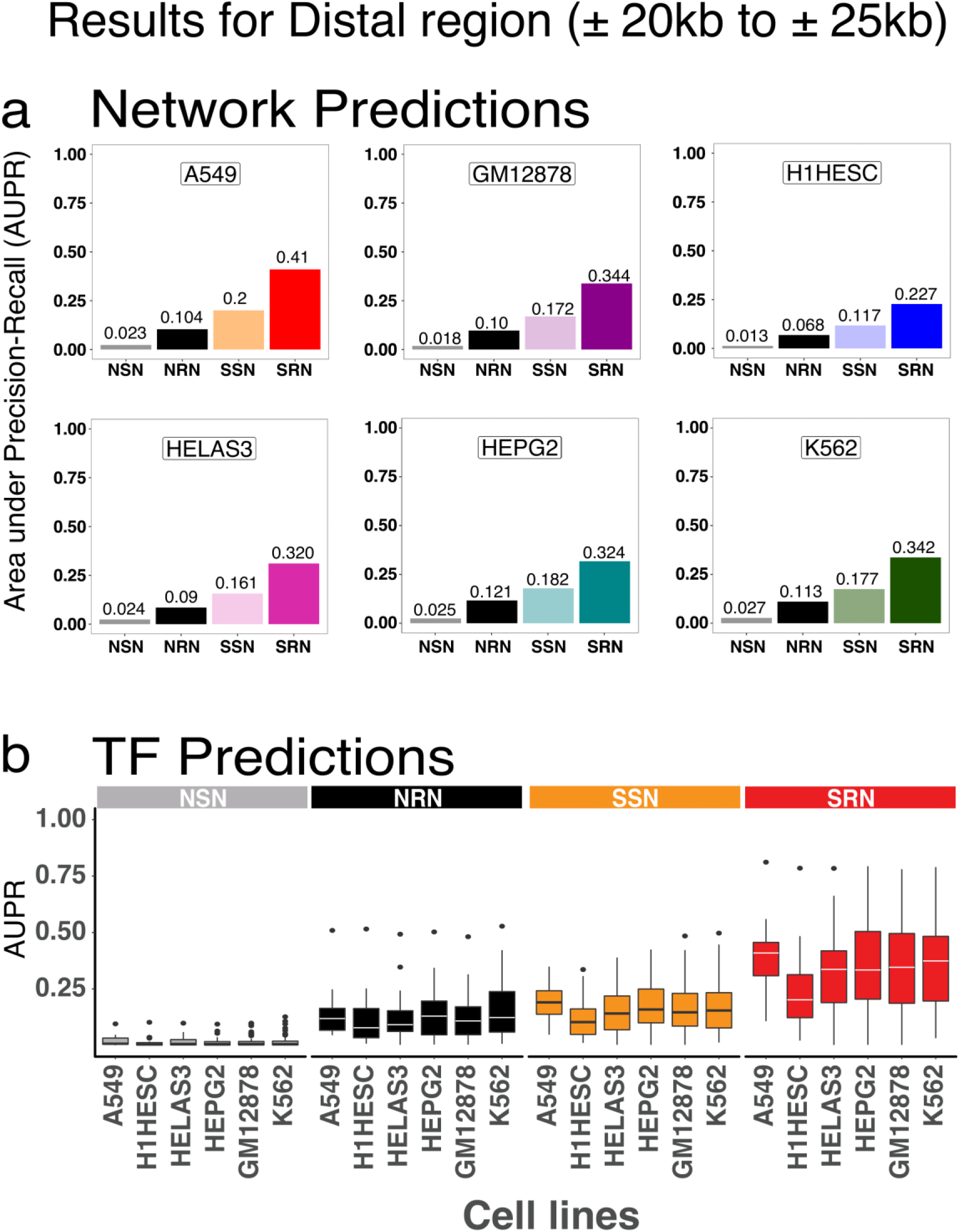
AUPR predictions for the distal region. **a.** AUPR scores for four types of regulatory models in six cell-lines. **b.** Distribution of AUPR scores across individual TF predictions. See also **Figure 4**.

**Supplemental Figure 4:**
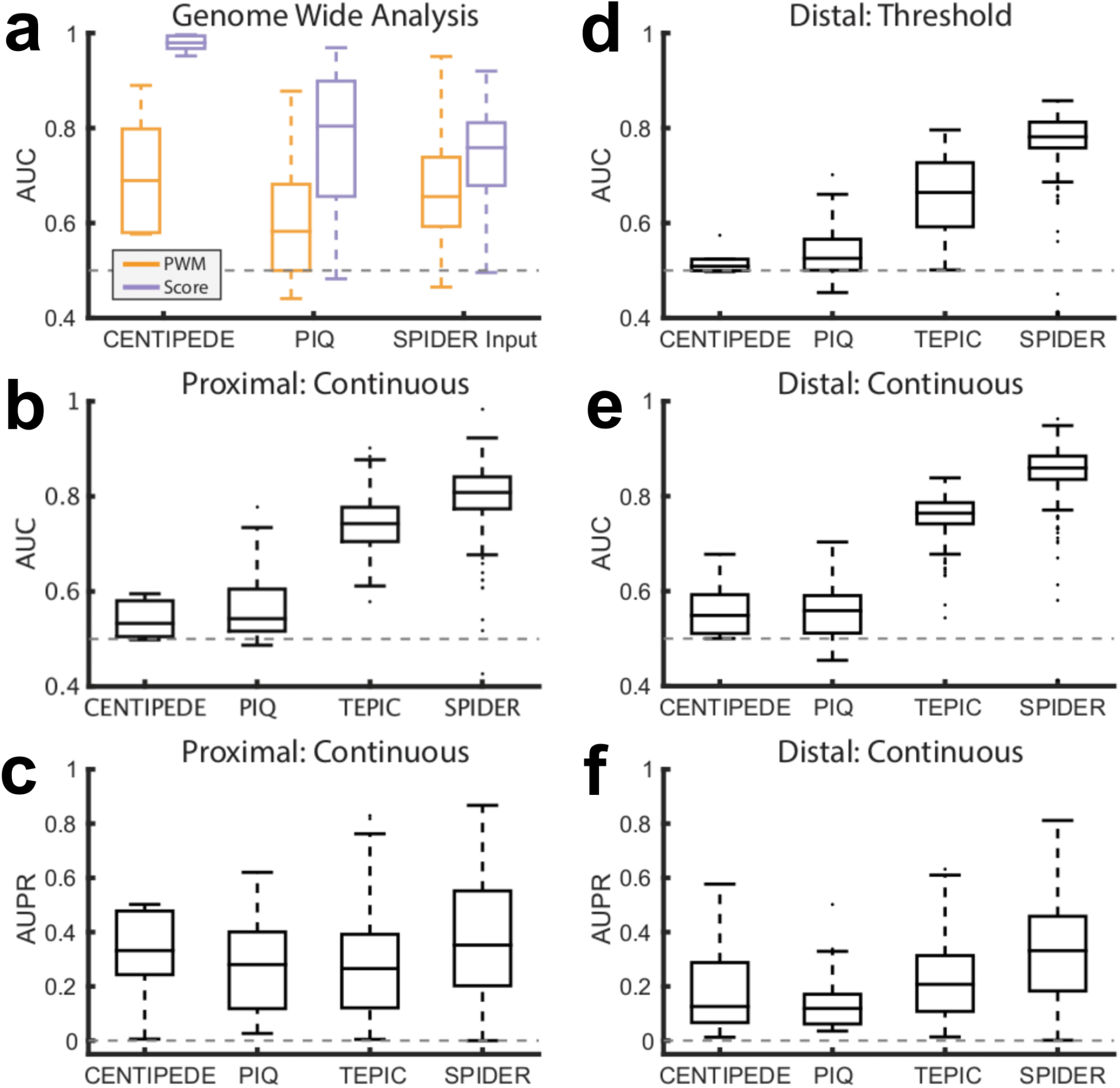
Detailed comparative benchmarking of SPIDER with other methods. **a.** Performance of CENTIPEDE, PIQ, and the input to SPIDER (the overlap of DNase peaks with motif predictions), when all TF motif locations across the genome are considered in the evaluation, as measured by AUC. **b-c**. Performance of the networks derived from the output of CENTIPEDE and PIQ, as well as the networks predicted by TEPIC and SPIDER, based on continuous (as opposed to thresholded) edge-weights, as measured by (**b**) AUC and (**c**) AUPR. **d-f**. Performance of the networks derived from the output of CENTIPEDE and PIQ, as well as the networks predicted by TEPIC and SPIDER, using distal regulatory regions, based on (**d**) thresholded and (**e-f**) continuous edge-weights, and evaluated using the (**d-e**) AUC and (**f**) AUPR. These analyses demonstrate that even though some methods, such as CENTIPEDE and PIQ, perform well at the genome-wide level (as shown in panel **a**), this does not necessarily translate into accurate network-level predictions (as demonstrated in panels **b-f**). See also **Supplemental Section S1**, **Supplemental Figure 1**, and **Figure 5**.

